# Spatial heterogeneity shapes microbial eco-evolutionary dynamics of soil carbon

**DOI:** 10.64898/2026.09.03.749071

**Authors:** Agathe Chave-Lucas, Régis Ferrière

## Abstract

The study of reciprocal influences between ecological and evolutionary processes has advanced considerably, yet integration between evolutionary biology and ecosystem-level ecology remains limited. Here we contribute to this integration by advancing the theory of eco-evolutionary feedbacks between soil microbial adaptation and soil-atmosphere carbon fluxes in a warming climate. We develop a spatially structured model of soil organic matter decomposition that represents microbial populations in microsites embedded in a bulk-soil matrix and focuses on exoenzyme production as a key resource-acquisition trait. The evolutionarily adapted investment in exoenzyme production is shaped by opposing selective forces: negative selection within microsites, where lower-investing mutants exploit exoenzymes as public goods, and positive selection in the soil matrix, where exoenzyme production directly benefits individual cells. Microsite density emerges as a critical determinant of microbial adaptation to warming and its consequences for soil carbon loss. Even small changes in microsite density across a threshold can reverse the ecosystem-level effect of adaptation, from buffering to amplifying carbon loss. High microsite density generally promotes buffering, whereas low microsite density has little effect in cool ecosystems but can strongly amplify carbon loss in warm ecosystems, especially when microbial mobility is low. These results identify soil spatial structure at microsite scale as a key mediator of microbial evolutionary adaptation and soil carbon-climate feedback under global environmental change, with implications for quantitatively improving Earth system models.

## Introduction

Soil ecological and biogeochemical processes play a major role in the Earth global carbon cycle. The soil carbon stock, in the form of organic and inorganic carbon, is about twice the atmospheric C pool (Scharlemann et al. 2014; Ontl and Schulte 2012). As a consequence, even a small fraction of soil carbon released to the atmosphere can have a large impact on the planet’s future climate. Soil organic carbon originates from dead organic matter decomposed by macro- and micro-organisms (Singh et al. 2010). Macro-organisms, such as worms, fragment large organic debris into smaller pieces, which are then further decomposed and consumed by microorganisms, resulting in metabolic products such as carbon dioxide and methane. As a result, soil micro-organisms, through their rate of organic matter decomposition, have a strong influence on greenhouse gas fluxes from soil to the atmosphere (Raich and Schlesinger 1992, D. S. Schimel 1995).

Soil microbial populations, with their large sizes and high levels of genetic variation, have the potential to evolve rapidly in response to environmental changes (Padfield et al. 2015, Schaum et al. 2017). However, how microbial adaptive traits change and how microbial adaptive trait evolution affects ecosystem processes such as organic carbon decomposition and CO_2_ respiration, remain poorly known (Classen et al. 2015, Abs, Leman, and Ferrière 2020, Abs, Chase, et al. 2024, Greenblum 2024). To advance our understanding of the adaptive variation of microbial traits and the consequences for ecosystem function, a useful framework is the Yield, Acquisition, Stress tolerance (Y-A-S) triangle rooted in life-history theory (Malik et al. 2020). This framework emphasizes tradeoffs between investing in growth efficiency, in the cellular machinery of resource acquisition, and in stress coping mechanisms. Given such tradeoffs, the conventional expectation is that under harsh environmental conditions, larger resource allocation into microbial traits pertaining to the stress-tolerant (S) strategy will be selected; in resource-rich environments, allocation to traits controlling yield (Y) may prevail; in competitive and resource-poor environments, allocation to traits pertaining to the acquisition (A) strategy should be favored (Malik et al. 2020).

In microbial populations experiencing similar stress levels, the Yield-Acquisition tradeoff is expected to be the primary axis of phenotypic variation. Heterotrophic microbes acquire their resources from outside the cell by secreting the extracellular enzymes (exoenzymes) on their surface or in their surroundings, thus driving organic matter decomposition. Consequently, a key determinant of the position of a population along the Yield-Acquisition axis is the individual investment in exoenzymes (Ramin and Steven D. Allison 2019). The production of exoenzymes, which has direct effects on the amount of substrate locally available for uptake, as well as the diffusive loss, both impose a substantial growth cost (Steven D. Allison et al. 2011, Scott et al. 2010, Smith and Chapman 2010, Lipson 2015, Traving et al. 2015, Wutzler et al. 2017). Furthermore, exoenzymes are public goods, which benefit even those cells that do not produce them. As a consequence, exoenzyme-producing strains are vulnerable to invasion by ‘cheaters’, i.e. genetically distinct individuals that invests less in exoenzyme production than the ‘resident’ strain (Steven D. Allison 2005, S. D. Allison 2012, Kaiser et al. 2015). In idealized well-mixed populations, strong negative selection should thus lead to minimal investment in exoenzyme production (Abs, Leman, and Ferrière 2020, Abs, Saleska, et al. 2025).

The spatial structure of microbial populations is expected to play an important role in the evolution and maintenance of cooperation traits such as exoenzyme production (Özkaya et al. 2017; Abs, Leman, and Ferrière 2020, Bonner et al. 2022). Yet the spatial heterogeneity of microbial populations and their environment has not been fully integrated in our understanding of microbial adaptation (Nunan, Schmidt, and Raynaud 2020). The case of biofilms, where the diffusion of products acting as public goods is limited, has been extensively studied (Özkaya et al. 2017). But biofilms only offer a small window into what happens in real soils, which can be described as complex, three-dimensional mixture of solid material and water-filled and gas-filled pores, where most of the microbial activity occurs in small groups clustering together in aggregates (Paul and Clark 1996, Ettema and Wardle 2002, Young and Crawford 2004, Nunan 2017). Aggregates create a mosaic of microsites which are embedded in a matrix of bulk soil, where the eco-physiological conditions are unfavorable and where cells may be isolated for prolonged periods of time (Wilpiszeski et al. 2019). In addition, microbes can move through the soil, with microbial mobility potentially playing an important role in soil function (Mason-Jones et al. 2025). At the scale of microsites, microbial mobility is most likely driven by passive transport (Abu-Ashour et al. 1994), with active motility contrained by high costs outside the wet range of matric potentials (Dechesne et al. 2010). Altogether, this raises the questions of how soil structure and microbial mobility (related to transport in and out of microsites) influence the evolution of exoenzyme production; what the ecological consequences are for ecosystem function such as decomposition and respiration; and how these consequences change as macroscopic soil parameters vary.

Here we develop a mathematical modeling framework that bridges the ecological effects of soil heterogeneity and microbial mobility and the evolution of exoenzyme production. The ecoevolutionary dynamics model captures the environmental feedbacks that operate at the local microbial population scale. The model predicts ecosystem-level impacts of eco-evolutionary responses to environmental change. Our analysis focuses on two environmental drivers, soil warming and variation in organic carbon supply, and evaluates the ecosystem-level effect of evolutionary adaptation on the soil-atmosphere carbon feedback.

## Methods

Soil microorganisms form metapopulations of microsites surrounded by a matrix of bulk soil (Ettema and Wardle 2002, Wilpiszeski et al. 2019, Fig. 1). Microsites support soil microaggregates that concentrate both water and microorganisms and where most of the microbial growth takes place in a complex three-dimensional network of pores (Nunan, Wu, et al. 2002, Raynaud and Nunan 2014, Wilpiszeski et al. 2019). Similarly to most microbial-explicit soil decomposition models, we lump prokaryotic decomposers and fungi into a single pool for a more parsimonious model parameterization (Crowther et al. 2019). Fungi are primarily found in copiotrophic environments (litter layers and rhizosphere, Frostegård and Bååth 1996). In these environments, and others where there are sufficient resources to facilitate hyphal spread, fungi would be expected to have a homogenizing or spatial averaging effect. This would occur because of their ability to overcome the spatial separation between resources that might arise in spatially heterogeneous environments. It has also been suggested that fungi can transport bacterial cells (known as the ‘fungal highway’, Kohlmeier et al. 2005), which would influence how bacterial decomposers explore heterogeneous space (Nunan, Schmidt, and Raynaud 2020). For these reasons, our model is better suited for oligotrophic environments dominated by bacterial decomposers where the influence of the plant rhizosphere is limited.

**Figure 1.**
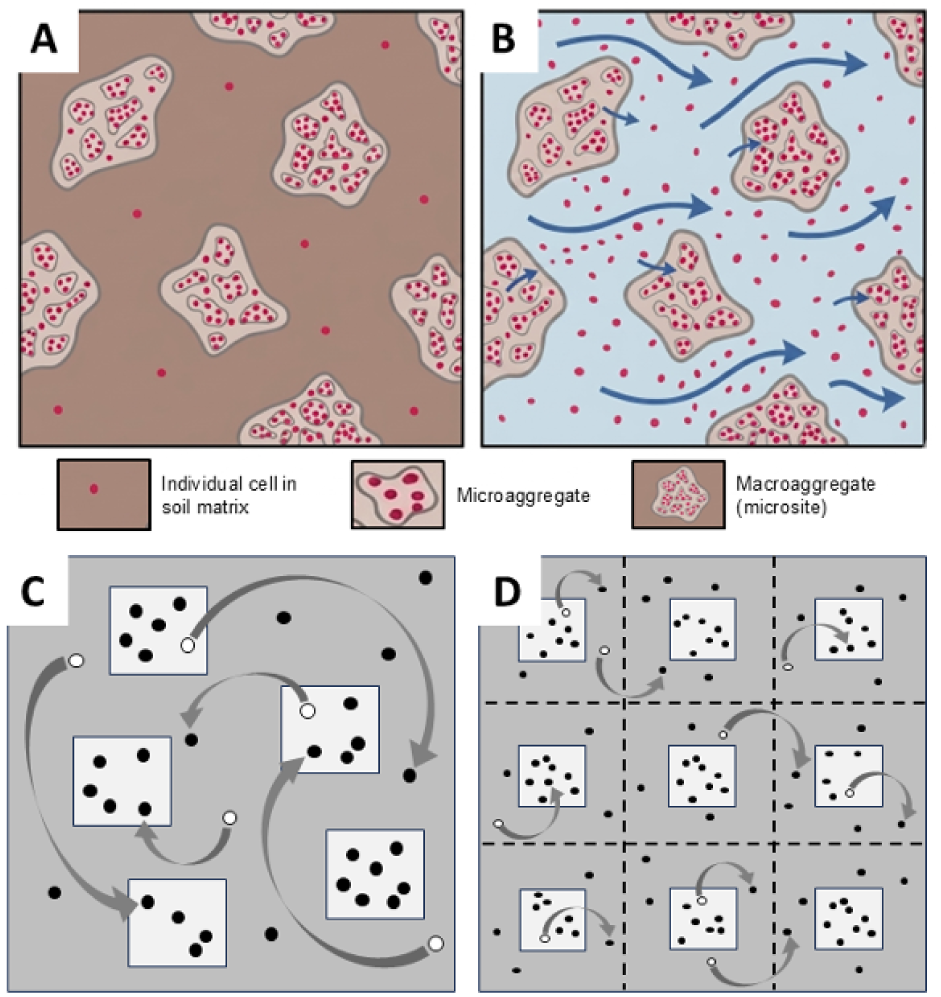
| Spatial structure of the soil microbial population. Soils can generally be viewed as complex three-dimensional structures consisting of packed aggregates in a matrix. Aggregates comprise clusters of mineral particles and organic carbon in which strong forces hold the particles together and make the structures persist through wetting events and mechanical disturbances of the soil. The networks of particles and cavities forming aggregates are connected during wetting events, which create variable flows of water and nutrients that can be accessed by soil microorganisms (Wilpiszeski et al. 2019). Small microaggregates (less than 250 *µ*m) assemble into larger macroaggregates (0.25 to 2 mm) with which most (about 90%) soil bacteria are associated (Ranjard et al. 2000). These macroaggregates thus provide favorable patches of bacterial habitat that we call microsites. The shape, distribution, organic matter content and water flow through and around the microsites form the base unit of structure-function relationships in most soil ecosystems. (A) Under drier conditions (brown matrix), microbial mobility events are rare and microbial density in the soil matrix surrounding microsites is low. (B) Under wetter conditions (blue matrix), water transport (and possibly active motility) can move bacteria in and out of microsites (blue arrows). (C, D) Cell dispersal models. Cell movement are indicated with arrows from initial location (open circle) to final location (filled circle). (A, B) adapted from Fig. 2 in Wilpiszeski et al. 2019. (C) Global dispersal model. Microsites (light gray) are randomly distributed in the matrix (dark gray). Cell movement is from microsite to matrix and from matrix to microsite, with effective movement between microsites being random and independent of the microsite spatial arrangement. (D) Local dispersal model. The soil representative volume is divided (dashed lines) into adjacent patches of matrix (dark gray), each containing one microsite (light gray). Cells can move from microsite to matrix inside a given patch, from matrix to matrix between adjacent patches, and from matrix to microsite inside a given patch. Our mathematical analysis assumes directional movement across the entire volume (e.g. from left to right in the figure), which could be due to directional water flow. This allows to collapse the model spatial dimension to one.

Our model assumes that microsites share similar physical features, as in J. Tang and Riley 2019; and microbes can move between microsites and the soil matrix, a possibility that Tang and Riley’s model ignores. Concentrations of substrates, exoenzymes and microbes differ within and between microsites, leading to a spatially structured soil system at mm scale. Thus, our investigation focuses on mobility and differential growth between microsites and the matrix, at mm scale; the complex spatial structure that exists within microsites at *µ*m scale, involving heterogeneous distributions of water, nutrients, and microbes in a network of pores of variable sizes (Chenu, Pouteau, and Nunan 2025 and references therein), will be addressed in future work (see Ebrahimi and Dani Or 2016 for an ecological modeling foundation).

To model the evolution of exoenzyme production, we use the adaptive dynamic framework, thus extending other recent applications (Abs, Leman, and Ferrière 2020, Abs, Saleska, et al. 2025) to the case of spatially heterogenous soil microbial populations. In this section, we build on Tang and Riley’s ecological model to include the mobility of microbes and their distribution across microsites and the soil matrix. We then derive invasion fitness for the spatially structured soil system and use invasion fitness to predict (i) the adapted state of the population to given environmental conditions; (ii) how the adapted state changes as the environmental conditions change; and (iii) the ecological consequences on ecosystem function, focusing on respiration and soil carbon loss.

We developed two versions of the ecological model, with global versus local dispersal (see Supplementary Materials), reflecting two endmember scenarios in a hierarchy of soil aggregates (Edwards and Bremner 1967, Oades and Waters 1991). In the global dispersal model, microorganisms and substrates may move among microsites via the soil matrix acting as a global pool; dispersal from one microsite to another is thus agnostic of the spatial location of microsites. In the local dispersal model, microorganisms and substrates only travel within their habitat patch, defined as a microsite and its immediate surroundings in the matrix, and between adjacent patches. These two end-member models yield similar ecological equations at equilibrium, similar forms of invasion fitness, hence similar eco-evolutionary outcomes (see Supplementary Materials).

### A spatially structured model of soil microbial exoenzyme production

To represent the dynamics of organic carbon in soil microsites, we use a CDMZ-type ecological model (Abs and Ferrière 2020, Manzoni and J. P. Schimel 2024) in which microbial biomass (*N_m_*) and the concentration of exoenzymes produced by microorganisms (*Z_m_*) are represented explicitly. Exoenzymes turn soil organic carbon (*C_m_*) into dissolved organic carbon (*D_m_*) which is available for uptake by the microorganisms. The ecological dynamics are governed by a system of four ordinary differential equations

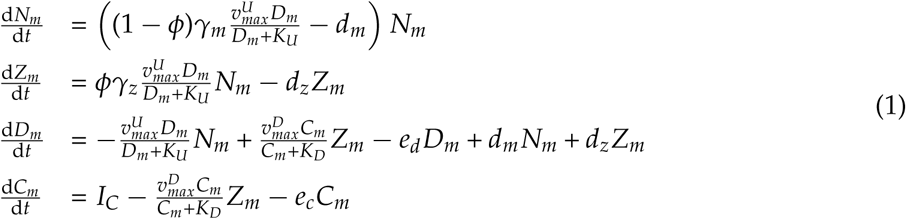

where *γ_m_*denotes the microbial growth efficiency, *γ_z_*measures enzyme production efficiency, *d_m_* is the microbial death rate and *d_z_* is the exoenzymes’ denaturation rate. The growth of microorganisms in microsites, the production of exoenzymes, and the production of dissolved organic carbon are given by Michaelis-Menten functions in which *v_max_* and *K* generically denote the maximum reaction rate and the half-saturation constant. Parameters *e_c_*and *e_d_* denote the leaching rates of, respectively, soil organic carbon and dissolved organic carbon, and *I_C_* is the input rate of soil organic matter in the soil.

The fraction of uptaken dissolved organic carbon that is invested in exoenzyme production is denoted by *ϕ*. Each microbial cell thus invests a fraction *ϕ* of uptaken carbon into exoenzyme production, and the remainder, (1 *− ϕ*), into growth, assuming that the minimal amount of carbon needed for cell maintenance is always available. The individual trait *ϕ* thus captures the fundamental tradeoff between acquisition and yield. The adaptive evolution of *ϕ* is constrained by this tradeoff and shaped by selection resulting from competition for the ‘public good’ of exoenzymes decomposing organic carbon.

To extend the model to the microbial metapopulation of microsites and bulk soil, we assume that the concentrations of enzymes and dissolved organic carbon are negligible in the soil matrix (in line with J. Tang and Riley 2019 and with empirical support from Leitner et al. 2017, Gao and DeLuca 2020 and Baveye and Aso 2021). Microorganisms may occur in the bulk soil at low but non-negligible densities (Raynaud and Nunan 2014), and they only have access to dissolved organic carbon resulting from the activity of their own exoenzymes. Mechanically, besides the relative isolation of individual cells, this may be further ensured when exoenzymes are attached to the membrane surface or by the production of extra-cellular polymeric substances (EPS), which can limit the diffusional loss of both the exoenzymes and the products of enzymatic decomposition (D. Or et al. 2007) (at a cost which would then be compounded, in our model, with the cost of exoenzyme production (Jayathilake et al. 2017)).

At the single-cell level, the microbial biomass in the bulk soil, denoted by *M*, is thus driven by

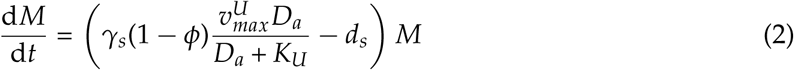

where *D_a_* denotes the expected concentration of dissolved organic carbon around an individual cell. The maximum uptake rate, 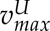, and half-saturation constant of microbial growth, *K_U_*, are assumed to be the same as in the microsites. The microbial growth efficiency, *γ_s_*, and death rate, *d_s_*, however, are different in the bulk soil. We assume that *γ_s_* is very small compared to *γ_m_*, reflecting the fact that the bulk soil matrix is a more hostile environment compared to microsites. The expected concentration, *D_a_*, of dissolved organic carbon resulting from the activity of exoenzymes produced by a single cell in the bulk soil and the expected concentration, *Z_a_*, of exoenzymes locally produced by a single cell, follows

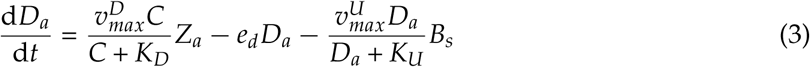

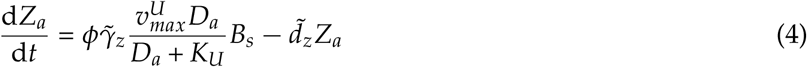

where *C* denotes the concentration of non-dissolved organic carbon in the bulk soil matrix, *B_s_*is the single-cell biomass concentration in the bulk soil, and parameters *γ̃*_*z*_ and *d̃_z_* represent the production efficiency and degradation rate of exoenzymes in the bulk soil (possibly different from the rates in microsites).

Assuming that the concentrations of dissolved carbon and exoenzymes equilibrate much faster than microbial biomass, we obtain the following equations for *Z_a_* and *D_a_* at equilibrium

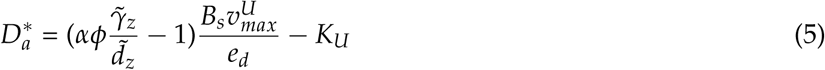

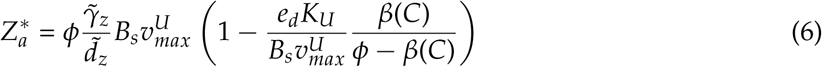

where

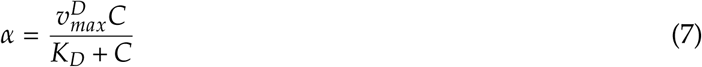

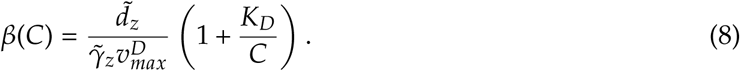

Combining equations (2) and (5) then yield the following equation for individual microbial biomass in the bulk soil

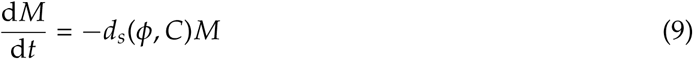

where *d_s_*(*ϕ*, *C*) is given by

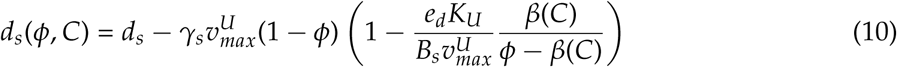

The results of our analysis qualitatively hold for the broad class of *d_s_* functions for which there exists a range of *ϕ* values (between 0 and 1) over which *d_s_* decreases as *ϕ* increases. Hence-forth we will be using the following notations

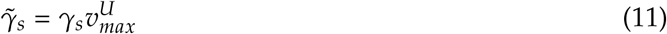

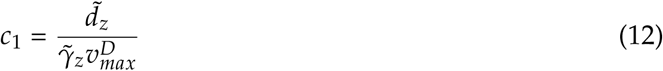

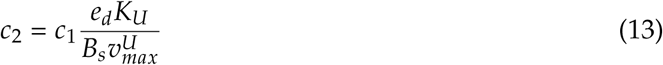

To obtain the two full ecological models describing microsites, bulk soil, and the fluxes between them, we use the single-cell biomass growth dynamics governed by Equation 9, and we assume that all microsites have identical properties (identical volumes and flux rates of substrate and microbes, as in J. Tang and Riley 2019) and that the microbial population in the bulk soil is sparse yet large, due to the significant extant of the soil matrix.

Even if the microsites have identical properties, they occur in different states in the full ecological models out of equilibrium (see Supplementary Materials). When a mutant appears in the system, the invaded microsites *de facto* vary in state. The two ecological models with multistate microsites are described in the Supplementary Materials, and both yield similar ecological and evolutionary results. In particular, they lead to similar invasion fitness functions. In the absence of mutants, assuming that all microsites are in the same state, the resident ecological model can be simplified into

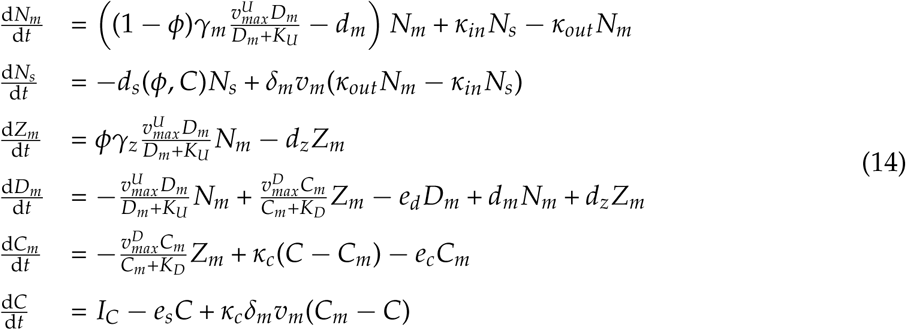

Here *N_s_* denotes microbial density in the bulk soil matrix, *C* is the concentration of non-dissolved organic carbon in the bulk soil matrix, and *I_C_* is the input of organic matter in the bulk soil matrix. Similarly to Tang and Riley’s model (J. Tang and Riley 2019), parameters *δ_m_* and *v_m_* measure the density and volume of microsites; *κ_c_* is the soil conductance of organic matter, i.e. the rate of carbon flow between microsites and the bulk soil matrix; *κ_in_* and *κ_out_* are conductance rates similarly defined for microbes: *κ_in_* is the flow rate of microbes from the bulk soil matrix to the microsites and *κ_out_*is the flow rate out of microsites. As microbes inside microsites are assumed to adhere to each other, we generally assume *κ_in_* to be larger than *κ_out_*.

At equilibrium, the soil carbon stock (non-dissolved) is measured by 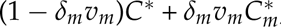, where *C^∗^* and 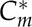 are the equilibrium concentrations of (non-dissolved) organic carbon in the bulk soil matrix and in the microsites, respectively. Solving the equation 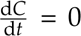 and using the fact that *e_S_* is very small compared to *κ_c_δ_m_v_m_*, we obtain:

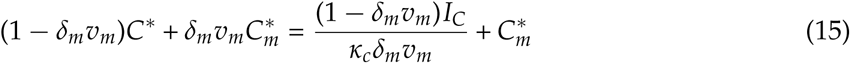

This shows that the response of the non-dissolved soil carbon to variation in environmental parameters (soil structure, carbon input, temperature) is mediated by an abiotic term [inlin e] and the biotically influenced term 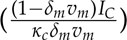 and the biotically influenced term 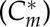. To properly evaluate the microbial ecological and evolutionary effects on the soil carbon stock, we focus our analysis on the response of soil carbon in microsites, 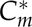, to environmental change.

### Invasion fitness in a spatially structured soil model

For a spatially structured soil model such as Equations 14, the calculation of invasion fitness for spatially homogeneous models (Parvinen and Seppänen 2016) does not apply straightforwardly. Here the calculation must take into account microbial population growth in microsites as well as in the bulk soil. Details of the calculation are given in the Supplementary Materials; hereafter we outline the main steps.

We focus on a mutant with trait *ϕ_mut_*arising in a resident population at ecological equilibrium. The mutant population is initially very small. Under these assumptions, the mutant population grows according to

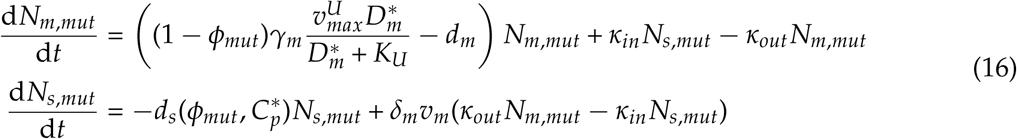

where 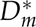 and 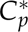 are, respectively, the equilibrium of dissolved organic carbon in the microsites and an estimate of non-dissolved organic carbon in the soil matrix for the resident population. Indeed, 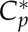 is assumed to be proportional to 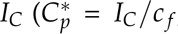, where *c _f_* is a constant), reflecting the assumption that most organic carbon enters the system as leaf litter. This simplification also allows us to reduce numerical instabilities and to explore the effect of other parameters more efficiently. The invasion fitness of the mutant trait *ϕ_mut_* in the resident population with trait *ϕ* is then given by

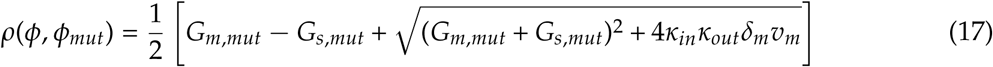

where

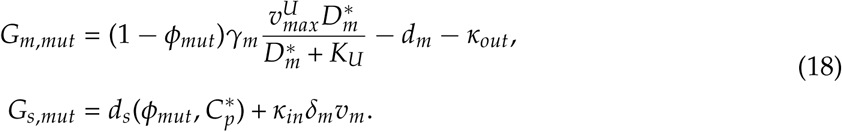

Potential end-points (attractors) or repellors of the evolutionary adaptive dynamics then obtain as zeroes of the selection gradient. Such evolutionary singularities, denoted by *ϕ^∗^*, are solutions of

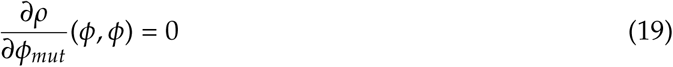

When there is no singularity (*i.e.* no solution to equation 19), selection is predicted to act directionally across the entire interval of feasible phenotypic values, in the direction predicted by the constant sign of the selection gradient.

### Ecological and evolutionary viability

Ecological viability is defined by Equation 14 possessing at least one locally stable equilibrium with non-zero microbial biomass. Evolutionary viability means that the eco-evolutionary dynamics themselves do not drive the microbial compartment to extinction (which would be an instance of ‘evolutionary suicide’, Ferrière and Legendre 2013). The condition for ecological viability is given by

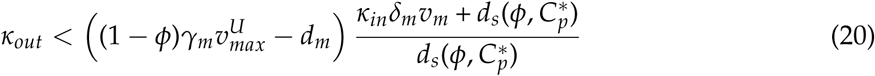

and the condition for evolutionary viability (*i.e.* no evolutionary suicide, given the ecological viability of the ancestral state) is given by

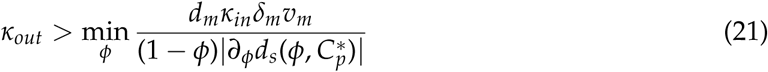

where *∂_ϕ_d_s_*is the first-order derivative of *d_s_* with respect to the first variable, *ϕ*.

If fluxes in and out of microsites are equal, *i.e. κ_in_* = *κ_out_* = *κ*, *κ* must remain between minimum and maximum bounds for the microbial population to be ecological viable. Specifically, if 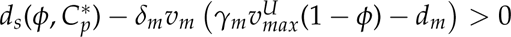, then *κ* is upper-bounded by

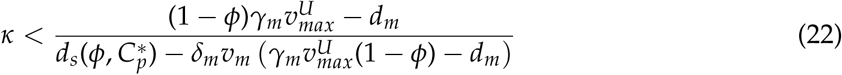

and lower-bounded according to

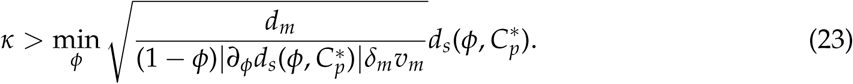

### Effect of temperature on microbial, enzymatic, and soil parameters

Temperature is assumed to affect the rates of within-cell enzymatic reactions driving uptake, assimilation, and biomass production. Temperature also affects the reaction rates of exoenzymes. As in Abs, Saleska, et al. 2025, the effects are captured by the Arrhenius model applied to the Michaelis-Menten parameters of resource acquisition and enzyme-driven decomposition (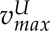, 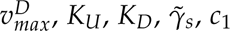, and *c*_2_). For example

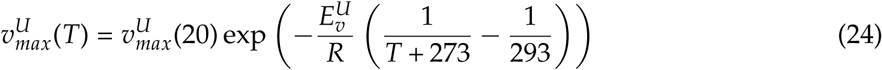

where *T* is measured in Celsius, 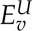 is the activation energy and *R* the universal gas constant. The values of 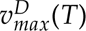, *K_U_*(*T*) and *K_D_*(*T*) are calculated using the activation energies 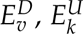 and 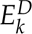, respectively. The temperature dependencies of *c*_1_ and *c*_2_ are given by

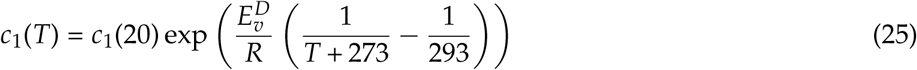

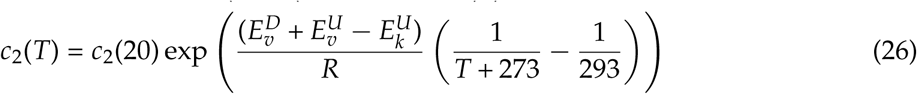

The microbial death rates, *d_m_* and *d_s_*, may either be constant and independent of temperature or follow an Arrhenius law as well, and both cases will be considered. The rates *d_m_* and *d_s_* are either temperature-independent, or have the same thermal dependency. Their activation energy is denoted by 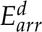. We assume that, at 30°C, the microbial death rates are equal, leading to the following relationships

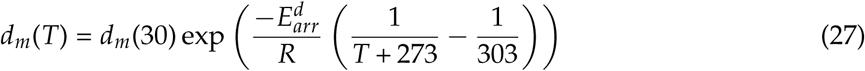

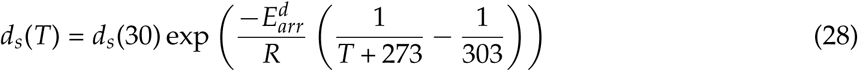

### Default parameter values

Default parameter values used in the Python code are listed in Table 1. The default parameter values for the microbial, enzymatic and organic and dissolved carbon parameters (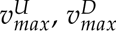, 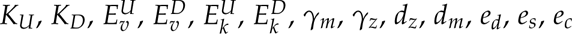 and *I_C_*) come from Abs, Saleska, et al. 2025. Approximate values for *δ_m_v_m_* and *κ_c_* come from J. Tang and Riley 2019, and other parameter values are specific to this study. *κ_in_* and *κ_out_* were constrained in such a way that the system is ecologically and evolutionary viable, *d_s_* was assumed slightly larger than *d_m_* (but of the same order of magnitude), *γ̃_s_* smaller than *d_s_*, so that microbes can never grow in the bulk soil, in accordance with J. Tang and Riley 2019 and Baveye and Aso 2021. Finally, *c*_1_ and *c*_2_ were constrained by taking reasonable values for *d̃_z_* and *γ̃_z_* (of the same order of magnitude as *d_z_* and *γ_z_*), and *c _f_* was constrained by assuming that the carbon available in the bulk soil matrix should be at least one order of magnitude smaller than if there were no microbes in the system.

**Table 1.**
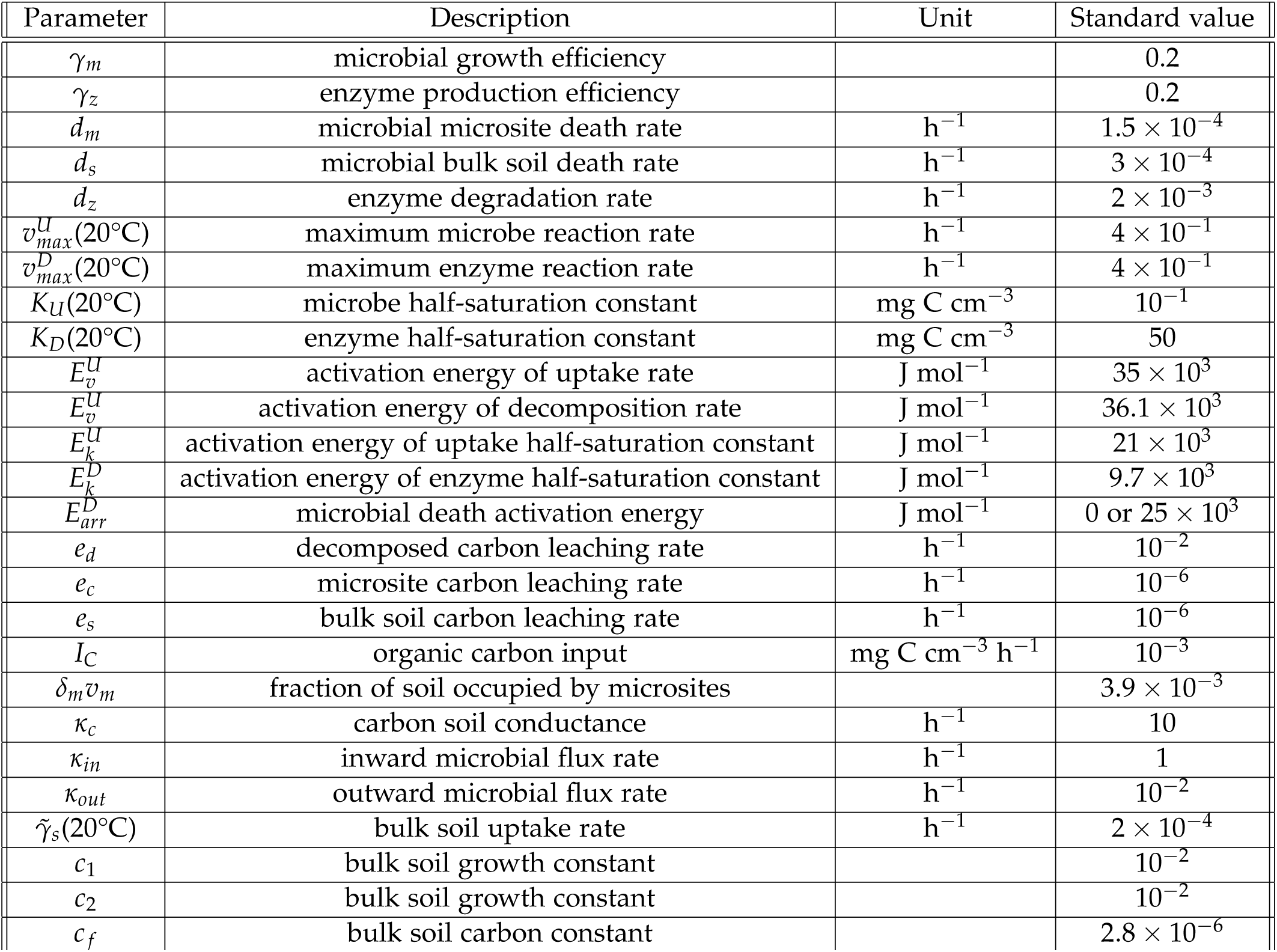
| Parameter list and default values.

## Results

To understand how microbial mobility and spatial population structure influence the system’s eco-evolutionary dynamics, first we address the ecological effects of microbial mobility and investment in exoenzyme, soil carbon input, and soil temperature on the system’s ecological viability and equilibrium soil carbon stock. Then we analyze the evolutionary viability of the system and the patterns of adapted investment in exoenzyme production in response to variation in microbial mobility, soil structure, and temperature. Finally, we quantify the effect of microbial evolutionary adaptation to warming on soil carbon depending on microbial mobility, soil structure, and exoenzyme parameters.

### Ecological dynamics

For given microbial, enzyme, and microsite parameters, ecological viability requires that the microsite outflux rate, *κ_out_*, be neither too small nor too large compared to the influx rate, *κ_in_* (eq. [20] and eq. [21]). The outside flux rate should not be too high since otherwise a large fraction of the population becomes exposed to the negative growth conditions of the bulk soil matrix, leading to ecological extinction. In particular, for a given value of *κ_in_*, there will be a maximum value of *κ_out_*to ensure ecological viability (Fig. S1A).

The investment in exoenzyme, *ϕ*, also plays a role in ecological viability: when *ϕ* reaches very low values, there is not enough decomposed carbon available for microbial uptake and growth; when *ϕ* reaches very high values, population viability is compromised by cells not investing sufficient energy in their own growth (Fig. S1A). Population extinction may also be driven by extreme values of temperature, carbon input (*I_C_*) and enzyme parameters (*γ_z_*, 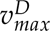 and *K_D_*) (results not shown).

In an ecologically viable system, the equilibrium soil carbon stock is smaller at higher temperature and for lower carbon input (*I_C_*) (Fig. 2). For a given carbon input and a given microbial investment in exoenzyme production, there is a temperature interval over which soil carbon decreases dramatically. For example, with carbon input *I_C_* = 10*^−^*^4^ mg C cm*^−^*^3^ h*^−^*^1^ and microbial decomposers characterized by *ϕ* = 0.05, soil carbon decreases more than three-fold as temperature rises from 15 °C to 25°C (Fig. 2A). With carbon input *I_C_* = 10*^−^*^3^ mg C cm*^−^*^3^ h*^−^*^1^ and microbial decomposers characterized by *ϕ* = 0.05, soil carbon decreases by one order of magnitude as temperature rises from 15°C to 18°C (Fig. 2B). The soil carbon stock is also sensitive to enzymatic parameters; in particular, soil carbon strongly decreases with enzyme conversion rates *γ_z_*(Fig. 2C) and strongly increases as the degradation rate of exoenzymes increases (Fig. 2D). In contrast, the ecological effect of microsite density, *δ_m_*, on the soil carbon stock is minimal (results not shown).

**Figure 2.**
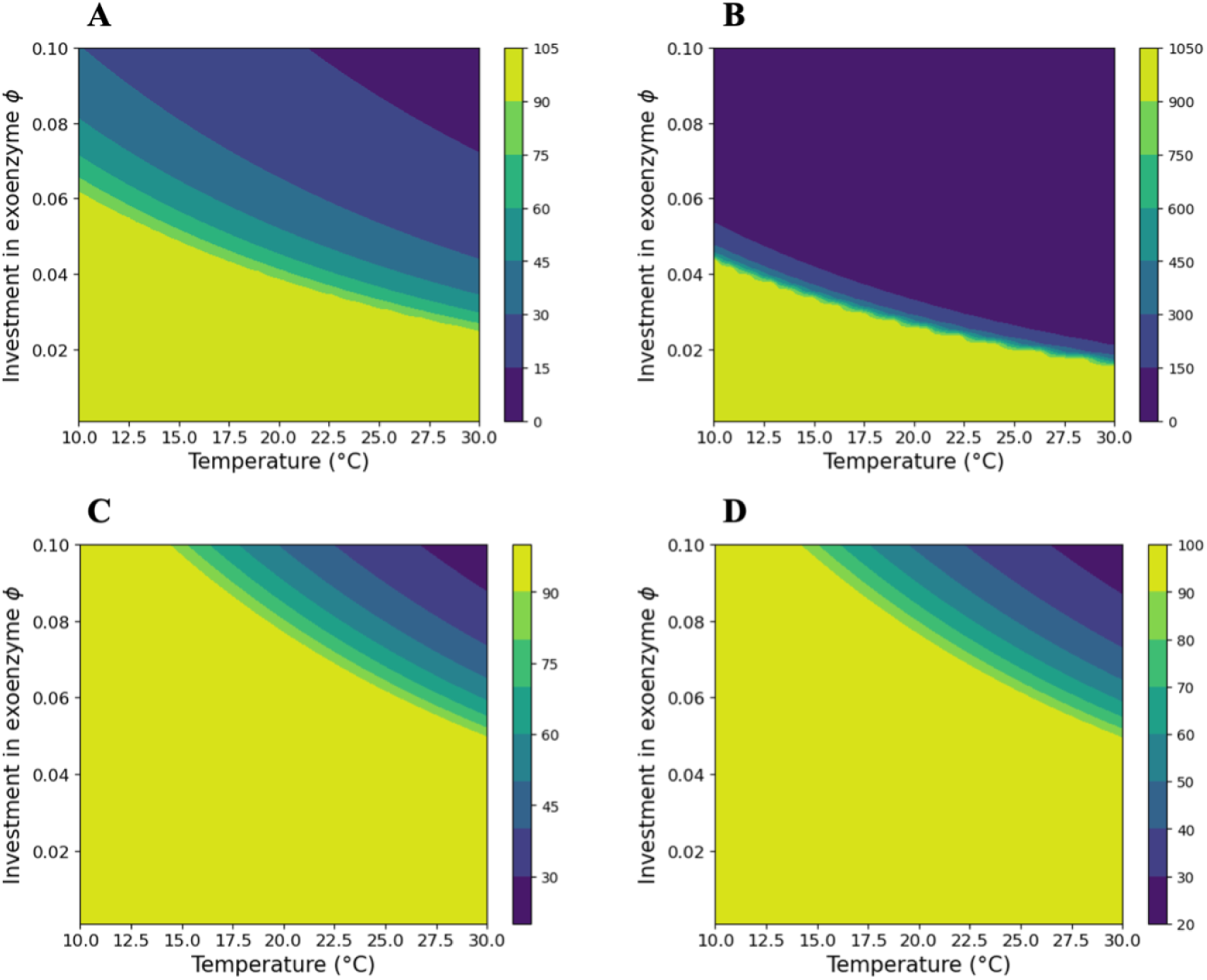
| Soil carbon stock in microsites (*C^∗^*) as a function of temperature and exoenzyme investment. (A) *I_C_* = 10*^−^*^4^ mg C cm*^−^*^3^ h*^−^*^1^. (B) *I_C_* = 10*^−^*^3^ mg C cm*^−^*^3^ h*^−^*^1^. (C) *I_C_* = 10*^−^*^4^ mg C cm*^−^*^3^ h*^−^*^1^ and *γ_z_* = 0.1. (D) *I_C_* = 10*^−^*^4^ mg C cm*^−^*^3^ h*^−^*^1^ and *d_z_* = 4 *×* 10*^−^*^3^ h*^−^*^1^. Other parameters are set to their default values (Table 1). Soil carbon stock is measured in mg C cm*^−^*^3^ (color code). The maximum values *C^∗^* = 100 mg C cm*^−^*^3^ in A, C and D, and *C^∗^* = 1000 mg C cm*^−^*^3^ in B, correspond to the balance of soil carbon input and leaching while the microbial population is extinct.

### Eco-evolutionary patterns of investment in exoenzyme production

The evolutionary viability of the system is strongly influenced by the microbial mobility parameters (Fig. S1B). When the microbial microsite outflux, *κ_out_*, is too small, there is no evolutionary singularity and evolutionary suicide occurs, whereby the population evolves towards self-extinction. For a given value of *κ_out_*, there is also a maximum value of *κ_in_*above which evolutionary suicide occurs. This is because, when the residence time of microbes inside microsites is too long compared to the bulk soil, cheating strains that invest less in exoenzymes systematically invade local populations within microsites; exoenzyme production is thus counterselected, leading to evolutionary suicide.

Under conditions preventing evolutionary suicide, the system evolves towards a viable singularity. This is the adapted value of the microbial investment in exoenzyme production, denoted by *ϕ^∗^*, which shows different patterns of variation with microbial mobility, microbial parameters, soil carbon input, and temperature depending on the density of microsites, *δ_m_* (Fig. 3 and Fig. S5). For any given combination of microbial mobility, soil carbon input, and temperature, there is a microsite density (*δ_m_* = 3 *×* 10^10^ in Fig. 3) at which *ϕ^∗^* is maximum. This means that for any given value of microbial mobility, soil carbon availability or temperature, evolutionary adaptation drives an increase in exoenzyme investment in response to an increase in microsite density in ecosystems with low microsite density (i.e. with *δ_m_* below 3 *×* 10^10^), as well as in response to a decrease in microsite density in ecosystems with high microsite density (i.e. with *δ_m_* above 3 *×* 10^10^). In addition, *ϕ^∗^* increases with microbial mobility at low microsite density whereas *ϕ^∗^* is essentially insensitive to microbial mobility at high microsite density (Fig. 3A). Irrespective of microsite density, *ϕ^∗^* decreases with soil carbon input (Fig. 3B). There is also a threshold on microsite density above which adaptation to warming drives exoenzyme investment to decrease and below which adaptation to warming drives exoenzyme investment to increase (Fig. 3C, D).

**Figure 3.**
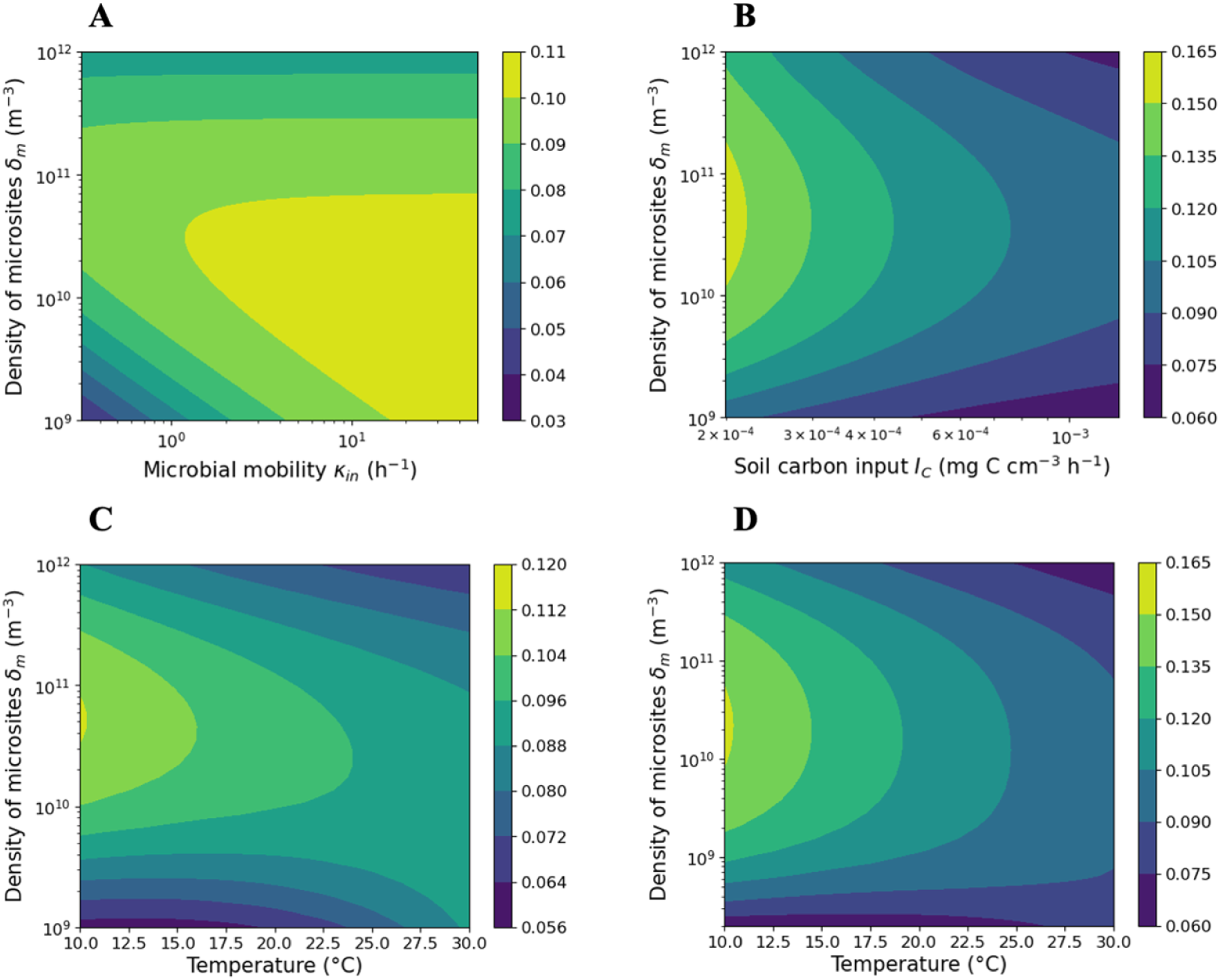
|Evolutionary adapted investment in exoenzyme production, *ϕ^∗^*, as a function of soil and microbial parameters. The adapted value is shown as a function of microsite density and (**A**) microbial mobility *κ_in_* (assuming *κ_out_* = 0.01*κ_in_*), (**B**) soil carbon input in the bulk soil, (**C**) soil temperature when microbial mortality is not temperature-dependent and (**D**) soil temperature when microbial mortality increases with temperature. In (**A**) and (**B**), temperature is fixed at 20 °C. Temperature-dependent mortality in (**D**) is such that the mortality rate at 30 °C is equal to the constant mortality rate in (**A**-**C**). Other parameters are set to their default values (Table 1).

The increase in adapted investment *ϕ^∗^* as mobility increases (Fig. 3A) reflects the fact that higher mobility results in a longer average residence time of microbes in the soil matrix, where mortality rises sharply when exoenzyme production is too low. Thus, exoenzyme investment represents a ‘private good’ in the bulk soil and individuals that invest more tend to be selected. As mobility increases, selection on the trait as a private good in the bulk soil tends to dominate selection against the trait as a public good in microsites, leading to a positive correlation between mobility and the adapted investment in exoenzyme production, *ϕ^∗^*.

The effect of mobility on the adapted exoenzyme production is modulated by the density of microsites (Fig. 3A). High microsite density facilitates microbes in the bulk soil reaching microsites, and this effect tends to dominate the influence of mobility on the adapted exoenzyme production. On the contrary, at low microsite density, variation in mobility has a strong effect on the adapted exoenzyme production. On the one hand, low mobility causes strong negative selection on exoenzyme production, since individual microbes tend to remain in their microsites where cheating strains are favored. On the other hand, high mobility combines with low microsite density to favor exoenzyme production as a private good. We note that this analysis uses *κ_in_*to measure microbial mobility, in a fixed ratio with *κ_out_*. The pattern of adapted exoenzyme production reported here is qualitatively insensitive to the value of this ratio (Fig. S2).

The effects of soil carbon input (Fig. 3B) and temperature (Fig. 3C) on the adapted investment in exoenzyme production also interact with microsite density. Indeed, for given soil carbon input or temperature, the adapted trait value is maximum at intermediate microsite densities. At high microsite densities, microbes tend to find new microsites in a shorter amount of time, which favors cheating strains; at low microsite densities, microbes will incur a high risk of death in the bulk soil, regardless of their exoenzyme investment, which weakens individual selection in the bulk soil and leads here again to the evolution of less cooperative strains. These effects are amplified under reduced organic carbon input (Fig. 3B), and this is robust to variation in temperature and microbial mobility, *κ_in_*(Fig. S3).

The pattern of adaptation to warming is more complex. At low microsite densities, the microbial population adapts to warming by investing more in exoenzyme production, whereas the opposite is true at high microsite densities (Fig. 3C). This pattern is qualitatively unaffected by variation in soil carbon input, *I_C_*, and microbial mobility, *κ_in_*(Fig. S4). When microbial mortality increases with temperature, the threshold microsite density at which the direction of the adaptive response to warming changes, is reduced by one order of magnitude (Fig. 3D).

### Eco-evolutionary response of soil carbon to warming

As microbial investment in exoenzyme production adapts to warming (Fig. 3C, D), ecological processes that depend on exoenzyme production, either directly (decomposition) or indirectly (resource acquisition, microbial growth), are impacted, with responses that potentially cascade up to the whole ecosystem level (soil carbon loss). Here we evaluate the hypothesis that the evolutionary adaptation of exoenzyme production to warming does impact soil carbon loss in a quantitatively significant manner. To this end, we compare the effect of warming with and without evolutionary adaptation. In scenarios with no evolutionary change, the investment in exoenzyme production is fixed at its adapted value at the baseline (initial) temperature. In scenarios with evolutionary adaptation, the investment in exoenzyme production continuously adapts along the temperature gradient.

Evolutionary adaptation of exoenzyme production to warming can have a substantial quantitative effect on soil carbon loss, either positive, hence amplifying the purely kinetic effect of warming on soil carbon loss; or negative, hence buffering the kinetic effect of warming on soil carbon loss. The direction of the effect strongly depends on the density of microsites (Figs. 4A, B vs. C, D) while being little sensitive to microbial mobility measured by microsite outflux, *κ_out_* (Figs. 4E, F, and Fig. S2). In the previous subsection, we highlighted the existence of a threshold on the density of microsites (about 4 *×* 10^9^ in Fig. 3) across which the adapted investment in exoenzyme production behaves differently in response to warming. The same threshold plays an important role in organizing the effect of adaptation to warming on soil carbon loss.

**Figure 4.**
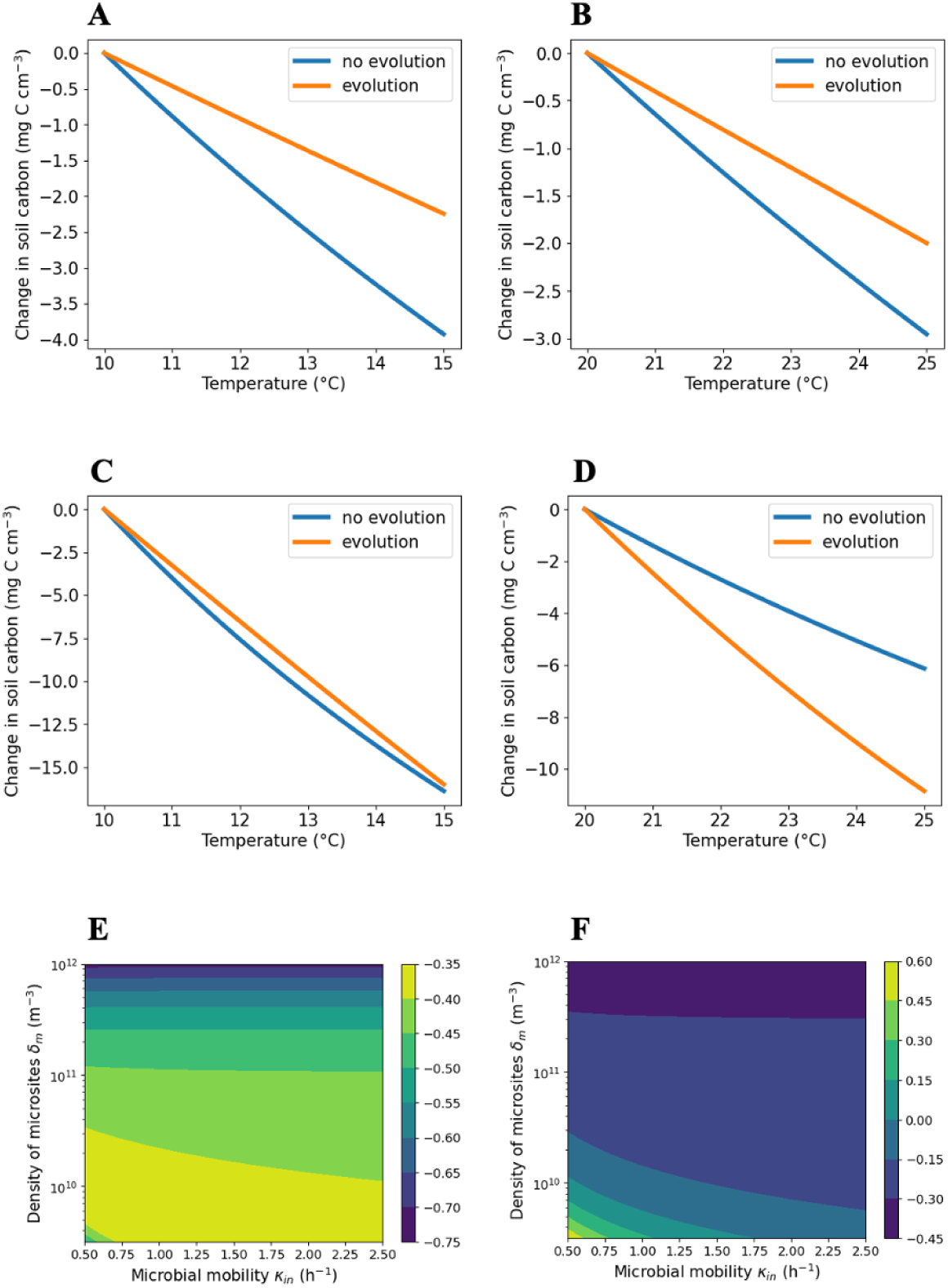
*|* Soil carbon loss with and without evolutionary adaptation of microbial exoenzyme production to warming: influence of spatial structure parameters when microbial mortality is not affected by temperature. (**A**) and (**B**) Microsite density *δ_m_* = 10^11^ m*^−^*^3^. (**C**) and (**D**) Microsite density *δ_m_* = 10^9^ m*^−^*^3^. (**E**) and (**F**): Difference in slopes between soil carbon loss without and with evolution for different values of microsite density and microbial mobility. When the difference in slopes is negative, then the soil carbon loss is larger without trait evolution, and vice versa. The ratio between *κ_out_* and *κ_in_* is kept constant, with *κ_out_* = 0.01*κ_in_*. (**E**) Difference in slopes between 10 and 15°C. (**F**) Difference in slopes between 20 and 25°C. Other parameters are set to their default values (Table 1).

When the microsite density exceeds the threshold, the adapted investment in exoenzyme production decreases as the temperature increases. As a result, the soil carbon stock changes (decreases) substantially less with evolutionary adaptation than without, both in cool (across the 10-15 °C range, Fig. 4A) and warm (across the 20-25 °C range, Fig. 4B) ecosystems. With microsite density below the threshold, soil carbon loss is generally much larger; for example four times larger when comparing the patterns of Figs. 4C and D to Figs. 4A and B. In this case, soil carbon loss with warming shows little difference with and without evolutionary adaptation in cool ecosystems (Figs. 4C), while soil carbon loss is dramatically amplified in warm ecosystems (two-fold in the numerical example of Figs. 4D). This difference comes from the fact that even though the adapted investment in exoenzyme increases with warming below the microsite density threshold, the increase is shallow across cool temperatures and more pronounced over warmer temperatures (Fig. 3C).

The same patterns qualitatively hold in systems where microbial mortality increases with temperature (figure S6), the main effect of the mortality temperature dependence being a lower threshold microsite density (Fig. 3D). Below the threshold, the adaptive increase of the investment in exoenzymes with temperature is much more shallow than in the case of temperature-independent mortality. Consequently, the amplifying effect of evolutionary adaptation to warming on soil carbon loss is reduced. Quantitatively, the buffering effect of microbial adaptation thus tends to dominate in both cool and warm ecosystems.

The soil carbon stock changes almost linearly with temperature across the +5 °C warming ranges considered here (Figs. 4A-D). The effect of evolutionary adaptation can thus be measured by the difference of the slopes of the soil carbon stock response to warming, with and without evolution, with the initial state being constrained at 10 °C in cool ecosystems and 25 °C in warm ecosystems (Figs. 4E, F). The slope difference becomes more negative (i.e. the buffering effect of evolutionary adaptation on soil carbon loss is enhanced) in soils with higher microsite density (in line with the comparison of Figs. 4A, B with Figs. 4C, D). The effect of microbial mobility is limited and more pronounced in warm ecosystems, where a reduction in mobility can turn the slope difference from negative to positive and drive large positive values, meaning that the amplifying effect of evolutionary adaptation on soil carbon loss is aggravated.

## Discussion

How microbial exoenzyme production varies in response to environmental change is key to predicting changes in local decomposition rates and how these changes scale up to influence soil respiration and feed back to atmospheric composition and climate. Our model shows that microbial mobility (which may be controlled largely by soil moisture) and investment in exoenzyme production both strongly influence the ecological viability of a simple carbon-focused, spatially-structured soil microbial ecosystem. Together with temperature and soil carbon input, and for given enzymatic parameters, they shape patterns of variation of the equilibrium soil carbon stock. In particular, increasing investment in exoenzyme production or temperature results in increased decomposition, hence a smaller soil carbon stock. In contrast, the soil carbon stock is little sensitive to the density of microsites.

For given environmental parameters (soil spatial structure, temperature, soil carbon input) and microbial mobility parameters, the evolutionarily adapted investment in exoenzyme production is determined by negative selection within microsites (where mutant strains that invest less than the resident strain are at a selective advantage) and positive selection in the soil matrix (where exoenzyme production provides a direct fitness benefit to individual cells). The adapted value tends to increase with increasing microbial mobility, decrease with increasing soil carbon input, and vary with increasing temperature depending on microsite density – decreasing if microsite density is above a threshold, increasing if microsite density is below the threshold – with a sensitivity to temperature that is higher in warmer environments.

These patterns of variation shape the impact of microbial adaptation to warming on soil carbon stock. In soils with relatively high microsite density, the adaptive response of exoenzyme production to warming tends to buffer soil carbon loss. In soils with relatively low microsite density, the effect of evolutionary adaptation on soil carbon loss is small in cool ecosystems while it can be large and amplifying in warm ecosystems. In the latter case, the combination of low microsite density and low microbial mobility further enhances the amplifying effect of evolutionary adaptation. Including a thermal dependence of microbial mortality in the model strengthens selection against exoenzyme production, which becomes less sensitive to changes in environmental temperature. As a consequence, the buffering effect of microbial adaptation on soil carbon loss is expected to occur across broader parameter ranges.

### Spatial structure shapes selection on exoenzyme production

Taking the evolutionarily adaptive potential of microbial traits into account (here, the individual investment in exoenzyme production, a key factor of resource acquisition) implies that these traits are not free parameters. Rather, given environmental parameters and fixed traits, adaptive traits are constrained to evolutionarily stable values, which are determined by the selection gradient acting on them. Selection on exoenzyme production is shaped by the population spatial stucture, with selection acting in opposite directions in microsites versus the bulk soil matrix. Within microsites, which are formed by complex three-dimensional networks of pores, exoenzymes diffuse and cheating strains that produce fewer or no exoenzymes can invade as they benefit from their neighbors’ exoenzymes while paying less of the cost of synthesizing them (or even no cost at all). This sets a negative selection component on the adaptive evolution of exoenzyme production. In contrast, in the bulk soil, individual cells acquire dissolved organic carbon resulting primarily from organic matter decomposed by their own exoenzymes; strains that invest more into exoenzyme are thus at a selective advantage. When the microbial mobility rates – especially *κ_out_*– increase, microbes spend more time in the bulk soil, which gives more weight to the positive selection component on exoenzyme production, hence selection of higher exoenzyme production.

Exoenzyme production thus evolves as a costly cooperation trait. Investment in this trait creates a “diffusible public good” (Driscoll and Pepper 2010) in microsites with high microbial density, which favors strains that invest less; and a “private good” in the bulk soil, where the microbial density is much lower and strains that invest more than others earn a direct fitness benefit. This “public-private good” combination was previously described by Driscoll, Hackett, and Ferrière 2016 as a possible mechanism to explain the evolutionary stability of exoenzyme production by toxic algae in aquatic ecosystems. Abs, Saleska, et al. 2025’s model of soil microbial exoenzyme production makes a similar assumption but without explicitly representing the spatial structure of the system from which the public and private goods emerge. In Abs, Leman, and Ferrière 2020, the evolution of exoenzyme production was addressed with a spatially explicit model, but with microsites forming a uniform coupled lattice; in that case, it was random local extinction and the spatial segregation of strains among microsites that made it possible for exoenzyme producers to resist invasion by cheating strains.

A large body of theory on diffusible public goods has shown that spatial structure can favor microbial cooperation by increasing assortment among producers and limiting access by cheaters. This logic appears in graph-based models of public-good diffusion, where cooperation is favored when diffusion is limited, colony dimensionality is low, and public-good decay is low (Allen et al. 2013); in individual-based models of digital microbes, where reduced cell diffusion or reduced public-good diffusion can each be sufficient to favor cooperation (Dobay et al. 2014); in models of “staying together,” where cooperator clusters evolve when they retain enough of the public goods they produce (Olejarz and Nowak 2014); and in the broader biofilm literature, where clonal segregation and local retention of benefits favor public-good production (Nadell et al. 2017). More recent work has extended this perspective by showing that directed migration can generate spatial patterns that promote cooperation (Funk and Hauert 2019), that microbial public-good cooperation depends strongly on life-cycle and spatial context (Cremer et al. 2019), and that spatial localization can preserve cooperative phenotypes in structured bacterial colonies (Monaco et al. 2022).

Most previous models, however, differ from ours in how they represent soil space and selection. Models of diffusible public goods in colonies, biofilms, lattices, graphs, continuous surfaces, or attached clusters generally assume that the same ecological rules operate everywhere. Spatial structure changes who interacts with whom and how far benefits diffuse, but not the qualitative nature of selection. Our model agrees with these studies at the scale of a microsite: exoenzyme production behaves as a diffusible public good, so diffusion and local mixing expose producers to cheating, whereas limited diffusion and producer clustering increase producer benefit. The key difference is that our model embeds this public-good phase within a larger soil life cycle. The novelty is therefore not spatial assortment alone, but the coexistence of public-good and private-good phases in the same microbial life cycle. Our findings substantiate Rillig, Muller, and Lehmann 2017 views on the role of soil aggregates as “evolutionary incubators”, implying that microsites and the soil matrix can contribute very different processes to the evolution of soil microbial communities.

This comparison also helps clarify which features of our results are general and which are soil-specific. The general feature is that exoenzyme production is favored when producers retain a large share of the benefits they create. This is the common thread linking graph models, digital microcolonies, biofilms, cooperator clusters, metapopulations, and decomposition models. The soil-specific feature is that benefit retention changes qualitatively across microenvironments. Within aggregates, benefit retention is reduced by diffusion and sharing; in the matrix, benefit retention is increased by isolation. Microsite density and microbial mobility therefore determine not simply the strength of assortment, but the time spent in two regimes of selection with opposite signs. This explains why increasing *κ_out_*favors higher exoenzyme production, why microsite density creates threshold behavior, and why soil structure can determine whether microbial adaptation buffers or amplifies soil carbon loss under warming.

More specifically, the decrease of the adapted value of exoenzyme production, *ϕ^∗^*, with soil carbon input does not depend on the functional form of microbial growth in the soil matrix – and neither does the high sensitivity of *ϕ^∗^* to the microsite density, *δ_m_*, enzyme efficiency cost, *γ_z_*, or enzyme degradation rate, *d_z_*, as these parameters do not appear in the bulk soil microbial growth rate. A specific form for microbial population growth in the bulk soil (Equation 2 and Methods) was derived and used in the model numerical simulations, but the results are largely insensitive to this functional form as long as exoenzyme production acts as a “private good” that is under positive selection in the bulk soil. In particular, the increase of *ϕ^∗^* with microbial mobility does not depend on the form of Equation 2; it is a general result emerging from the spatial structure of the population.

### Microbial mobility

Microbial mobility, quantified by the flux rates *κ_in_* and *κ_out_* in and out of microsites, encompasses all movements of microbes in the soil, either passive or active. To constrain these rates, we used reference values for *κ_m_*, the flow of substrate in and out of microsites (J. Tang and Riley 2019), and assumed that *κ_in_*and *κ_out_* were several orders of magnitude smaller than *κ_m_*, as microbial cells are expected to diffuse much less in and out of microsites than substrate molecules (Miyamoto and Shimono 2020) and show low motility in general (Dechesne et al. 2010). Similarly to *κ_m_*, the soil conductance of a substrate, being dependent on soil humidity and compaction, we expect *κ_in_* and *κ_out_* to increase with soil humidity and decrease with compaction (Abu-Ashour et al. 1994; Yang and van Elsas 2018; J. Tang and Riley 2019). Under wetter conditions, microbes (and substrates) diffuse and move more between microsites (Dechesne et al. 2010). On the contrary, in compact or dry soil, microbial mobility is even more limited. Empirical measurements of microbial mobility and its environmental factors are urgently needed, building on previous experiments such as those reported by Buffi et al. 2025, and in line with Mason-Jones et al. 2025 who recently called for the reassessment of active microbial mobility in soil and its functional consequences.

We also expect parameters *κ_in_* and *κ_out_* to depend on microbial parameters, including microbial size, adhesion in microsites, as well as other biotic features of the soil environment, such as the presence of other micro- and macro-organisms (Abu-Ashour et al. 1994; Kohlmeier et al. 2005). Our results suggest that adapted exoenzyme investment should be higher in aerated soils with a high density of transport agents such as hyphal fungi or mobile macro-organisms. This is consistent with observations showing that microbial exoenzyme production increases with the abundance of soil animal microfauna (Lipiec et al. 2016; Buivydaitė et al. 2023). By contrast, the adapted exoenzyme investment is expected to be much reduced in compact soils with little to no microfauna, such as intensive agricultural soils, which tend to be compacted by heavy machinery and where toxic chemical inputs can severely impact the microfauna (Maggi and F. H. Tang 2021). This aligns with empirical evidence showing that aerobic prokaryote respiration negatively correlates with soil compaction (Hartmann et al. 2013; Longepierre et al. 2021), although those observations could also reflect limited access to nutrients in compacted soils.

### Adaptive response of exoenzyme production to warming

Empirical studies have begun to document patterns of variation in exoenzyme production, with contrasted results. Some studies report exoenzyme production decreasing with elevated temperatures while others document increasing enzymatic activity and production with warming (Fanin et al. 2022; Daunoras, Kac_̌_ergius, and Gudiukaitė 2024). Our results support the expectation of a diversity of responses shaped by microbial adaptation and in which the soil spatial structure, as modeled here, plays an important role.

Indeed, a small variation in the density of microsites (across the threshold highlighted in Figs. 3 and 4) can change the magnitude and even revert the sign of the adaptive response of exoenzyme production to warming. In soil systems sparsely populated in microsites (below the threshold), the adapted trait, *ϕ^∗^*, is expected to increase with temperature. In soil systems with a high density of microsites (i.e. above the threshold), *ϕ^∗^* is predicted to decrease with soil temperature. This differs from Abs, Saleska, et al. 2025’s model, wherein *ϕ^∗^* was generally predicted to increase with warming. In their model, the public good of dissolved organic carbon decreases as temperature increases, due to faster uptake; strains that produce more exoenzymes are then at a selective advantage thanks to the marginal benefit of their private good. In Abs, Saleska, et al. 2025, spatial effects are limited to substrate diffusion and are modeled implicitly, with a private good being defined as a localized increase in DOC available to a cell that invests more in exoenzymes than the population average cell. This spatially-explicit model shows that separating out the effects of the public good (on growth in microsites) and private good (on survival in the soil matrix) can result in microsite density shaping the direction of the adaptive response of exoenzyme production to warming.

The threshold and pattern of variation of *ϕ^∗^* with temperature quantitatively depend on microbial mobility, carbon input as well as microbial mortality and growth rates. These key features of the soil system, together with microsite density, may co-vary with temperature it-self, either as a direct effect (e.g. physiological effect of temperature on microbial motility) or through correlated effects (e.g. physical effect of temperature on soil porosity, water film continuity or water transport) and concurrent changes in other environmental drivers (e.g. changes in precipitation affecting water distribution and material transport); all effects that may further vary across spatial and temporal scales (Oishy et al. 2025). In spite of the complexity of these effects and their interactions, two main scenarios can be proposed, based on the results shown in Fig. 3, to capture their qualitative impact on the adaptive response of exoenzyme production to warming, again depending on the range of microsite densities within which the system may vary.

In the first scenario, warming correlates with wetter conditions, which may result in higher microsite density, higher microbial mobility, and larger soil carbon input. In the second scenario, warming correlates with drier conditions, which is assumed to correlate with lower microsite density, lower microbial mobility, and smaller soil carbon input. A strong positive adaptive response of exoenzyme production to warming, i.e. a strong increase of *ϕ^∗^* with temperature, is expected in the first, warmer-wetter scenario applied to soils evolving in a range of relatively low microsite density, and in the second, warmer-drier scenario applied to soils evolving in a range of relatively high microsite density. In contrast, a strong negative adaptive response of exoenzyme production to warming, i.e. a strong decrease of *ϕ^∗^* with temperature, is expected in the warmer-wetter scenario applied to soils evolving in a range of relatively high microsite density, and in the warmer-drier scenario applied to soils evolving in a range of relatively low microsite density. In the next subsection, we revisit these responses to predict their consequences in terms of change in soil carbon stock.

### Microbial adaptation and soil carbon loss

Temperature and exoenzyme production have direct effects on soil carbon stock (Fig. 2): increasing temperature and/or exoenzyme concentration accelerates decomposition and increase microbial respiration, resulting in a smaller soil carbon stock. Conditions under which evolutionary adaptation to warming drives exoenzyme production up are therefore expected to amplify soil carbon loss. Our analysis identifies microsite density as a critical parameter determining whether microbial evolutionary adaptation drives such amplification. More precisely, warm ecosystems with relatively low microsite density are especially prone to adaptation-driven amplification of soil carbon loss.

This result extends the previous findings of Abs, Saleska, et al. 2025 by showing that adaptation-driven amplification of soil carbon loss is not limited to cold ecosystems. Moreover, adaptation-driven buffering of soil carbon loss is generally expected in both cool and warm ecosystems characterized by relatively high microsite density. Our discussion of climate change scenarios, where warming correlates with changes in precipitation that jointly impact soil environmental and microbial spatial structure, predicts large amplification in response to wetter (respectively drier) conditions affecting soils with relatively low (respectively high) microsite density; and large buffering in response to wetter (respectively drier) conditions affecting soils with relatively high (respectively low) microsite density. It is worth noting that the influence of microsite density, *δ_m_*, is mediated in the model by the fraction of soil occupied by microsites, *δ_m_v_m_*. Some studies have already highlighted the important effect of pore size (directly related to *v_m_*) for microbial activity and carbon use efficiency (Chenu, Pouteau, and Nunan 2025). Such studies start filling the longstanding void of empirical knowledge about the spatial distribution and activity of microogranisms in soils and their functional impact on soil carbon (Baveye, Otten, et al. 2018). In our model, the strong influence of *δ_m_v_m_*calls for more empirical assessment of this compound parameter and its variation across space and time.

Soil carbon loss is also sensitive to the biochemical cost of the specific exoenzymes produced by the microbial community, measured by parameter *γ_z_*, and the enzyme degradation rate, *d_z_*. These effects stem from the direct influence that both parameters have on the exoenzyme concentration. Even though the singularity *ϕ^∗^* does not depend on *γ_z_* or *d_z_*, lower values of *γ_z_* or higher values of *d_z_* can revert the direction of the effect of evolutionary adaptation to warming on soil carbon loss, from positive to negative. As enzymatic parameter values may vary with the type of enzyme, as well as with temperature and other environmental factors (Baker and Steven D. Allison 2017; Alves et al. 2021; Sousa et al. 2023), more detailed models of enzymatic activity are warranted to refine our predictions.

## Conclusion

The study of the reciprocal influence between ecological and evolutionary processes has advanced considerably over the last two decades. Yet the integration of evolutionary biology and ecosystem ecology remains limited. Here we contribute to this integration by advancing the theory of eco-evolutionary feedbacks between soil microbial adaptation and soil-atmosphere carbon fluxes in a warming climate. A simple representation of soil structure (microsites and matrix) uncovers the importance of microbial distribution and mobility in shaping these feedbacks. The density of microsites emerges as a key determinant of the adaptive response of microbial exoenzyme production to warming and its impact on soil carbon loss. All else being equal, even a small change of microsite density across a critical threshold can revert the direction of the effect of microbial adaptation on soil carbon loss, between buffering and amplifying. Thus, models that ignore microbial eco-evolutionary feedbacks in soil carbon decomposition may substantially underestimate or overestimate soil carbon loss in response to climate warming, depending on spatially-related soil and microbial characteristics.

Our results emphasize the need for advancing empirical measurements of microbial mobility and the spatial distribution of microsites. Temporal fluctuations in these parameters could be the norm, with mobility responding to wet and dry periods, and microsite density fluctuating seasonally with carbon input. With such environmental variability, microbial adaptive stress responses such as dormancy are expected. Advancing our theory to capture both spatial and temporal variation of relevant soil and microbial characteristics and the joint evolution of resource acquisition and stress resistance will be the next step towards the integration of soil microbial eco-evolutionary processes into a predictive framework aimed at improving Earth system models and projections of carbon cycling and climate.

## Supporting information

Supplementary Materials (Equations and Figures)

## Acknowledgments

A.C.-L. is supported by the French Ministry of Agriculture for her doctoral research. R.F. is supported by the Biology Integration Institute EMERGE, U.S. National Science Foundation (NSF) Biology Integration Institutes Program, award 2022070, and by the NSF Growing Convergence Research, award 2121155. We thank Elsa Abs, Antonin Affholder and Scott Saleska for discussion.

## Statement of Authorship

A.C.-L. and R.F. conceptualized the study. A.C.-L. developed the model, did the math and coding, and generated the results. A.C.-L. and R.F. interpreted the results. A.C.-L. wrote the first draft of the manuscript, which was revised by R.F. Both authors finalized the manuscript.

## Data and Code Availability

The code for this work, which generates all figures in the Results and Supplementary Materials, is available on a Figshare Repository.

The link to the repository is: https://figshare.com/s/68fb5adcf256faa256c9

## Notes

### Competing Interest Statement

The authors have declared no competing interest.

