## Supplementary Materials (Equations and Figures) for "Spatial heterogeneity shapes microbial eco-evolutionary dynamics of soil carbon"

### 1 Ecological models

For simplicity and parsimonious parametrization, we treat the community of soil bacterial decomposers as a single functional group. Cells grow in microsites and can move (primarily through transport by water or other organisms) in and out of microsites and through the bulk soil matrix. Here we develop two ecological models: a global dispersal model, which ignores the spatial arrangement of microsites with respect to each other, and a local dispersal model, in which microbial movement is limited to neighboring microsites and interstitial matrix. Hereafter we show that at equilibrium, both models yield similar ecological and evolutionary outcomes.

As a starting point we use a simple and general resource-consumer model for the dynamics of microbial biomass, $M$, and substrate abundance, $S$, measured in unit of carbon concentration

$$\begin{aligned}\frac{\mathrm{d}M}{\mathrm{d}t} &= f_e(S)M - dM \\ \frac{\mathrm{d}S}{\mathrm{d}t} &= I_e - \tilde{f}_e(S)M\end{aligned}$$

where $f_e(S)$ is the growth rate of the microbial group in the environment $e$, $\tilde{f}_e(S)$ is the microbial consumption rate of the substrate in environment $e$, $I_e$ is the substrate input in that environment, and $d$ is the microbial death rate. Different forms of $f_e$ and $\tilde{f}_e$, in different types of environment $e$, will be considered.

#### 1.1 Global dispersal

Here we are considering a volume $V$ of soil in which a large number $n$ of microsites are randomly distributed. The volume of each microsite is denoted by $v_m$ (identical across microsites). The microsite volume $v_m$ is assumed to be very small compared to $V$. Substrates and microbes can diffuse from a microsite to the bulk soil, and vice versa, but not directly from one microsite to another. The soil matrix is modeled as one compartment, and we assume that the concentration of both substrate and microbes is homogeneous across the soil matrix.

In the following each microsite is indexed by its identification number $i$ (between 1 and $n$). Then $S_i$ and $N_i$ denote the substrate concentration and microbial abundance (both measured in unit of carbon mass per unit volume) in microsite $i$, respectively. The production of substrate in microsite $i$ is denoted by $p_i$. The substrate and microbial concentrations in the bulk soil matrix are denoted by $S$ and $N_s$, respectively.

##### Substrate concentrations in the microsites and bulk soil matrix

Similarly to Tang and Riley 2019's model, the time variation of substrate concentration in microsite $i$ is given by

$$\frac{\mathrm{d}S_i}{\mathrm{d}t} = -\tilde{f}_i(S_i)N_i + p_i + \kappa_m(S - S_i)$$

where $\kappa_m$ is the soil conductance as defined by Tang and Riley 2019, and $\tilde{f}_i$ the microbial consumption rate in microsite $i$. The time variation of substrate concentration in the bulk soil is given by

$$\frac{\mathrm{d}S}{\mathrm{d}t} = I_S - D_S S - \tilde{f}_s(S)N_s + \frac{\kappa_m v_m}{V}\sum_{i=1}^{n}(S_i - S)$$

where $I_S$ is the input of substrate in the soil matrix, and $D_S$ is the diffusion rate towards the exterior of the system (e.g. the atmosphere).

##### Microbial concentrations in the microsites and soil matrix

Likewise, the time variation of microbial concentration in the microsites and soil matrix are given by

$$\frac{\mathrm{d}N_i}{\mathrm{d}t} = f_i(S_i)N_i - d_i N_i + \kappa_{mic}(N_s - N_i)$$

$$\frac{\mathrm{d}N_s}{\mathrm{d}t} = f_s(S)N_s - d_s N_s + \frac{\kappa_{mic} v_m}{V}\sum_{i=1}^{n}(N_i - N_s)$$

where $\kappa_{mic}$ is the analogue of soil conductance for the microbes. In order to keep the algebra simple, and without loss of generality, the following equations and proofs are written for $\kappa_{in} = \kappa_{out} = \kappa_{mic}$; the same algebra can be developed for values of $\kappa_{in}$ and $\kappa_{out}$ that are different, only to be more tedious. This parameter (as addressed in the Discussion) is likely to be influenced by the type of soil, the size and other physical characteristics of the microsites, and on adhesion and active motility of microbes.

**Identical microsites**

If all microsites are identical, then all $p_i$'s assume a common value $p$, and likewise for the $f_i$'s $(= f)$, $\tilde{f}_i$'s $(= \tilde{f})$, and $d_i$'s $(= d)$. Then all $S_i$'s and $N_i$'s are equal, to $S_m$ and $N_m$, respectively, and governed by

$$\begin{aligned} \frac{\mathrm{d}S}{\mathrm{d}t} &= I_s - D_S S - \tilde{f}_s(S)N_s + \kappa_m \delta_m v_m (S_m - S) \\ \frac{\mathrm{d}N_s}{\mathrm{d}t} &= f_s(S)N_s - d_s N_s + \kappa_{mic} \delta_m v_m (N_m - N_s) \end{aligned}$$

where $\delta_m$ is the density of microsites, as in Tang and Riley 2019, i.e. $\delta_m = \frac{n}{V}$.

#### 1.2 Local dispersal

In this model, the soil is described as a lattice of patches, each containing one microsite. We assume that cells and substrates can move in and out between any given microsite and its surrounding patch; and between adjacent patches. Thus, movement between two neighboring microsites is a three-step process: moving out of one microsite into the surrounding patch matrix, then into the adjacent patch matrix, and finally into the microsite located in that patch. For the sake of mathematical tractability, we only treat the highly simplified case of a one-dimensional array of patches. In spite of its simplicity, this model may still capture the dynamics of a portion of soil where some steady water flow imposes a one-dimensional axis of transport.

In this one-dimensional model, we consider a volume $V$ of soil subdivided into $n$ adjacent patches (with $n$ very large), each containing one microsite (volume $v_m$) surrounded by bulk soil. In the following mathematical derivation, patches and microsites are indexed by $i$. For any patch $i$ (between 2 and $n-1$), movement of cells and substrates occur within patch $i$ and between patch $i$ and patches $i-1$ and $i+1$. We keep using the notations $S_i$ for substrate concentrations, $p_i$ for substrate production, and $N_i$ for microbial concentration (in unit of carbon concentration) inside microsite $i$. In addition, we denote substrate and microbial concentrations in the matrix of patch $i$ by $S_{n+i}$ and $N_{n+i}$, respectively; and the substrate production rate in any patch matrix by $I_b$.

##### Substrate concentrations in the microsites and bulk soil matrix

As in Tang and Riley 2019's model, the substrate concentration in microsite $i$ varies in time according to

$$\frac{\mathrm{d}S_i}{\mathrm{d}t} = -\tilde{f}_i(S_i)N_i + p_i + \kappa_m(S_{n+i} - S_i)$$

where $\kappa_m$ is the soil conductance (as defined by Tang and Riley 2019) and $\tilde{f}_i$ is the microbial consumption rate in microsite $i$. The substrate concentration in the $i$ patch matrix varies according to

$$\frac{\mathrm{d}S_{n+i}}{\mathrm{d}t} = I_b - D_S S_{n+i} + D_a(S_{n+i-1} + S_{n+i+1} - 2S_{n+i}) - \tilde{f}_B(S_{n+i})N_{n+i} + \frac{n\kappa_m v_m}{V}(S_i - S_{n+i})$$

where $\tilde{f}_B$ is the microbial consumption function in the patch matrix (assumed to be constant across all patches), $D_S$ is the diffusion rate towards the exterior of the system (e.g. the atmosphere), and $D_a$, the substrate diffusion rate between adjacent patches.

##### Microbial concentrations in the microsites and bulk soil matrix

Likewise, the microbial concentration in microsite $i$ varies according to

$$\frac{\mathrm{d}N_i}{\mathrm{d}t} = f_i(S_i)N_i - d_i N_i + \kappa_{mic}(N_{n+i} - N_i)$$

where $\kappa_{mic}$ is the analogue of soil conductance for the cells. And the microbial concentration in the $i$ patch matrix varies according to

$$\frac{\mathrm{d}N_{n+i}}{\mathrm{d}t} = f_B(S_{n+i})N_{n+i} - d_B N_{n+i} + D_p(N_{n+i-1} + N_{n+i+1} - 2N_{n+i}) + \frac{n\kappa_{mic}v_m}{V}(N_i - N_{n+i})$$

where $f_B$ and $d_B$ are respectively the growth function and death rate in the bulk soil portions, and $D_p$ the microbial diffusion rate between adjacent bulk soil portions.

**Identical microsites**

With all microsites sharing the same characteristics, we have $p_i = p$, $f_i = f$, $\tilde{f}_i = \tilde{f}$ and $d_i = d$, for all $i$'s. Then the above equations can be simplified. In particular, if we assume periodic boundary conditions (i.e. the quantities of cells and substrates moving out of patch $n$ "to the right" are taken to constrain the quantities of cells and substrates moving into patch 1 "from the left"), then we have, in any patch $i$, $S_i = S_m$, $N_i = N_m$, $S_{n+i} = S$, and $N_{n+i} = N_s$. This leads to

$$\begin{aligned}
\frac{\mathrm{d}S}{\mathrm{d}t} &= I_b - D_S S - \tilde{f}_B(S)N_s + \kappa_m \delta_m v_m (S_m - S) \\
\frac{\mathrm{d}N_s}{\mathrm{d}t} &= f_B(S)N_s - d_B N_s + \kappa_{mic} \delta_m v_m (N_m - N_s)
\end{aligned}$$

where $\delta_m$ is the density of microsites, as in Tang and Riley 2019. Note that here $\delta_m = \frac{n}{V}$, since there is one microsite in each patch of volume $\frac{V}{n}$. We thus conclude that with identical microsites, and periodic boundary conditions, the mathematical form of the simplified models is identical bewteen local and global dispersal cases.

### 2 Invasion fitness

Here we derive the invasion fitness of a mutant strain that interacts with a resident strain at ecological equilibrium (Metz, Nisbet, and Geritz 1992). The mutant strain may differ from the resident strain in any trait, denoted generically by $x$, that affects the microbial growth rate, $f$. Therefore, we use the subscript *mut* to refer to the concentrations and growth rates of the mutant strain in microsites and in the soil matrix environment.

#### Global dispersal model

At ecological equilibrium, the concentrations of substrate in each microsite $i$ and in the soil matrix are denoted by $S_i^*$ and $S^*$, respectively. The growth of a rare mutant in microsite $i$ and in the soil matrix is governed by

$$\frac{\mathrm{d}N_{i,mut}}{\mathrm{d}t} = f_{i,mut}(S_i^*)N_{i,mut} - dN_{i,mut} + \kappa_{mic}(N_{s,mut} - N_{i,mut})$$

and

$$\frac{\mathrm{d}N_{s,mut}}{\mathrm{d}t} = f_{s,mut}(S^*)N_{s,mut} - d_s N_{s,mut} + \frac{\kappa_{mic} v_m}{V}\sum_{i=1}^{n}(N_{i,mut} - N_{s,mut})$$

Using the notation $a_i = f_{i,mut}(S_i^*) - d$ and $a_s = f_{s,mut}(S^*) - d_s$, we can rewrite this system of linear differential equations in a vector form

$$\begin{pmatrix} \frac{\mathrm{d}N_{1,mut}}{\mathrm{d}t} \\ \frac{\mathrm{d}N_{2,mut}}{\mathrm{d}t} \\ \vdots \\ \vdots \\ \frac{\mathrm{d}N_{n,mut}}{\mathrm{d}t} \\ \frac{\mathrm{d}N_{s,mut}}{\mathrm{d}t} \end{pmatrix} = \begin{pmatrix} a_1 - \kappa_{mic} & 0 & \cdots & \cdots & 0 & \kappa_{mic} \\ 0 & a_2 - \kappa_{mic} & 0 & \cdots & 0 & \kappa_{mic} \\ \vdots & 0 & \ddots & \ddots & \vdots & \vdots \\ \vdots & \vdots & \ddots & \ddots & 0 & \vdots \\ 0 & 0 & \cdots & 0 & a_n - \kappa_{mic} & \kappa_{mic} \\ \frac{\kappa_{mic} v_m}{V} & \frac{\kappa_{mic} v_m}{V} & \cdots & \cdots & \frac{\kappa_{mic} v_m}{V} & a_s - \frac{n\kappa_{mic} v_m}{V} \end{pmatrix} \begin{pmatrix} N_{1,mut} \\ N_{2,mut} \\ \vdots \\ \vdots \\ N_{n,mut} \\ N_{s,mut} \end{pmatrix}$$

Then the invasion fitness of the mutant is given by the largest eigenvalue of the above matrix, denoted by $M_{n+1}$. After some algebra, we find that the $n+1$ (real) eigenvalues of $M_{n+1}$ are the $n+1$ roots of the characteristic polynomial

$$P(X) = \left( (X - a_s + \frac{n\kappa_{mic} v_m}{V}) \prod_{i=1}^{n} (X - a_i + \kappa_{mic}) \right) - \frac{\kappa_{mic}^2 v_m}{V} \sum_{i=1}^{n} \left( \prod_{j=1, j\neq i}^{n} (X - a_j + \kappa_{mic}) \right).$$

Assuming that all microsites have identical properties, we have all $a_i$'s equal, and by denoting $a_i = a$, we can simplify the characteristic polynomial into

$$\begin{aligned} P(X) &= \left((X-a_s+\tfrac{n\kappa_{mic}v_m}{V})(X-a+\kappa_{mic})^n\right)-n\tfrac{\kappa_{mic}^2 v_m}{V}(X-a+\kappa_{mic})^{n-1} \\ &= (X-a+\kappa_{mic})^{n-1}\left((X-a_s+\tfrac{n\kappa_{mic}v_m}{V})(X-a+\kappa_{mic})-n\tfrac{\kappa_{mic}^2 v_m}{V}\right) \end{aligned}.$$

Using $\delta_m = \frac{n}{V}$, we obtain

$$P(X) = (X-a+\kappa_{mic})^{n-1}\left((X-a_s+\kappa_{mic}\delta_m v_m)(X-a+\kappa_{mic})-\kappa_{mic}^2\delta_m v_m\right).$$

The largest root is the invasion fitness, $s_x(x_{mut})$, hence

$$s_x(x_{mut}) = \frac{1}{2}\left[a-\kappa_{mic}+a_s-\kappa_{mic}\delta_m v_m+\sqrt{(a-\kappa_{mic}-a_s+\kappa_{mic}\delta_m v_m)^2+4\kappa_{mic}^2\delta_m v_m}\right].$$

#### Local dispersal model

At equilibrium, the concentrations of substrate in microsite $i$ and in the $i$ matrix patch are denoted by $S_i^*$ and $S_{n+i}^*$, respectively. Then the mutant population growth in microsite $i$ and in matrix patch $i$ obeys

$$\frac{\mathrm{d}N_{i,mut}}{\mathrm{d}t} = f_{i,mut}(S_i^*)N_{i,mut}-d_iN_{i,mut}+\kappa_{mic}(N_{n+i,mut}-N_{i,mut})$$

and

$$\begin{aligned} \tfrac{\mathrm{d}N_{n+i,mut}}{\mathrm{d}t} &= f_{s,mut}(S_{n+i}^*)N_{n+i,mut}-d_sN_{n+i,mut} \\ &+ D_p(N_{n+i-1,mut}+N_{n+i+1,mut}-2N_{n+i,mut})+\tfrac{n\kappa_{mic}v_m}{V}(N_{i,mut}-N_{n+i,mut}) \end{aligned}$$

Hereafter we use the notations $a_i = f_{i,mut}(S_i^*)-d_i$ and $b_i = f_{s,mut}(S_{n+i}^*)-d_s$ and we define the matrix $M_{2n}$ as

$$M_{2n} = \begin{pmatrix} A_n & B_n \\ C_n & D_n \end{pmatrix}$$

where

$$A_n = \begin{pmatrix} a_1 - \kappa_{mic} & 0 & \cdots & \cdots & 0 \\ 0 & a_2 - \kappa_{mic} & 0 & \cdots & 0 \\ \vdots & 0 & \ddots & \ddots & \vdots \\ \vdots & \vdots & \ddots & \ddots & 0 \\ 0 & 0 & \cdots & 0 & a_n - \kappa_{mic} \end{pmatrix}$$

$B_n = \kappa_{mic} I_n$, $C_n = \frac{n\kappa_{mic} v_m}{V} I_n$, and $D_n$ is the circulating matrix

$$D_n = \begin{pmatrix} b_1 - \frac{n\kappa_{mic} v_m}{V} - 2D_p & D_p & 0 & \cdots & 0 & D_p \\ D_p & b_2 - \frac{n\kappa_{mic} v_m}{V} - 2D_p & D_p & 0 & \cdots & 0 \\ 0 & D_p & \ddots & \ddots & \ddots & \vdots \\ \vdots & 0 & \ddots & \ddots & \ddots & 0 \\ 0 & \vdots & \ddots & \ddots & \ddots & D_p \\ D_p & 0 & \cdots & 0 & D_p & b_n - \frac{n\kappa_{mic} v_m}{V} - 2D_p \end{pmatrix}$$

Then the system of linear differential equations which governs the mutant population growth from low density can be rewritten as

$$\begin{pmatrix} \frac{\mathrm{d}N_{1,mut}}{\mathrm{d}t} \\ \frac{\mathrm{d}N_{2,mut}}{\mathrm{d}t} \\ \vdots \\ \vdots \\ \frac{\mathrm{d}N_{n,mut}}{\mathrm{d}t} \\ \frac{\mathrm{d}N_{n+1,mut}}{\mathrm{d}t} \\ \frac{\mathrm{d}N_{n+2,mut}}{\mathrm{d}t} \\ \vdots \\ \vdots \\ \frac{\mathrm{d}N_{2n,mut}}{\mathrm{d}t} \end{pmatrix} = M_{2n} \begin{pmatrix} N_{1,mut} \\ N_{2,mut} \\ \vdots \\ \vdots \\ N_{n,mut} \\ N_{n+1,mut} \\ N_{n+2,mut} \\ \vdots \\ \vdots \\ N_{2n,mut} \end{pmatrix}$$

The mutant invasion fitness is given by the largest (real) eigenvalue of the matrix $M_{2n}$ or, equivalently, by the largest root of the caracteristic polynomial $\det(XI_{2n})$. To compute that largest root, first we note that $\det(XI_{2n} - M_{2n}) = \det\left((XI_n - A_n)(XI_n - D_n) - B_n C_n\right)$. Then we assume the case where all patches have identical properties, which implies that all $a_i$'s and all $b_i$'s are equal, i.e. $a_i = a$ and $b_i = b$. Then we have $A_n = (a - \kappa_{mic})I_n$ and $(XI_n - A_n)(XI_n - D_n) - B_nC_n = (X - a + \kappa_{mic})(XI_n - D_n) - \frac{n\kappa_{mic}^2 v_m}{V} I_n$. The matrix $(XI_n - A_n)(XI_n - D_n) - B_nC_n$ is a circulating matrix, whose determinant can be explicitly calculated. We use a classical result on circulating matrices, according to which, if

$$F_n = \begin{pmatrix} \alpha_0 & \alpha_1 & 0 & \cdots & 0 & \alpha_1 \\ \alpha_1 & \alpha_0 & \alpha_1 & 0 & \cdots & 0 \\ 0 & \alpha_1 & \ddots & \ddots & \ddots & \vdots \\ \vdots & 0 & \ddots & \ddots & \ddots & 0 \\ 0 & \vdots & \ddots & \ddots & \ddots & \alpha_1 \\ \alpha_1 & 0 & \cdots & 0 & \alpha_1 & \alpha_0 \end{pmatrix}$$

then

$$\det(F_n) = \prod_{k=0}^{n-1} \left( \alpha_0 + 2\alpha_1 \cos\left(\frac{2k\pi}{n}\right) \right).$$

Applying this result we obtain

$$\begin{aligned} &\det(XI_{2n} - M_{2n}) \\ &= \prod_{k=0}^{n-1} \left( (X - a + \kappa_{mic})(X - b + \frac{n\kappa_{mic}v_m}{V} + 2D_p) - \frac{n\kappa_{mic}^2 v_m}{V} - 2D_p \cos\left(\frac{2k\pi}{n}\right)(X - a + \kappa_{mic}) \right) \\ &= \prod_{k=0}^{n-1} \left( (X - a + \kappa_{mic}) \left( X - b + \frac{n\kappa_{mic}v_m}{V} + 2D_p(1 - \cos\left(\frac{2k\pi}{n}\right)) \right) - \frac{n\kappa_{mic}^2 v_m}{V} \right). \end{aligned}$$

The $2n$ eigenvalues $(\lambda_i)_{0 \le k \le 2n-1}$ of $M_{2n}$ are the $2n$ roots of the polynomial $\det(XI_{2n} - M_{2n})$. With the notation $g_k = b - \frac{n\kappa_{mic}v_m}{V} + 2D_p(\cos\left(\frac{2k\pi}{n}\right) - 1)$ for any $k$ between 0 and $n-1$, the eigenvalues can then be expressed as

$$\lambda_{2k} = \frac{1}{2} \left[ (a - \kappa_{mic}) + g_k + \sqrt{(a - \kappa_{mic} - g_k)^2 + \frac{4n\kappa_{mic}^2 v_m}{V}} \right],$$

$$\lambda_{2k+1} = \frac{1}{2} \left[ (a - \kappa_{mic}) + g_k - \sqrt{(a - \kappa_{mic} - g_k)^2 + \frac{4n\kappa_{mic}^2 v_m}{V}} \right].$$

The eigenvalue $\lambda_{2k}$ is largest when $g_k$ is largest, that is when $g_k = g_0 = b - \frac{n\kappa_{mic}v_m}{V}$. The mutant invasion fitness is thus given by

$$s_x(x_{mut}) = \frac{1}{2}\left[(a-\kappa_{mic})+b-\frac{n\kappa_{mic}v_m}{V}+\sqrt{\left(a-\kappa_{mic}-b+\frac{n\kappa_{mic}v_m}{V}\right)^2+\frac{4n\kappa_{mic}^2v_m}{V}}\right].$$

In this model, $\frac{n\kappa_{mic}v_m}{V} = \kappa_{mic}\delta_m v_m$, and therefore the same mathematical expression is recovered for the mutant invasion fitness in both global and local dispersal scenarios.

### Supplementary figures:

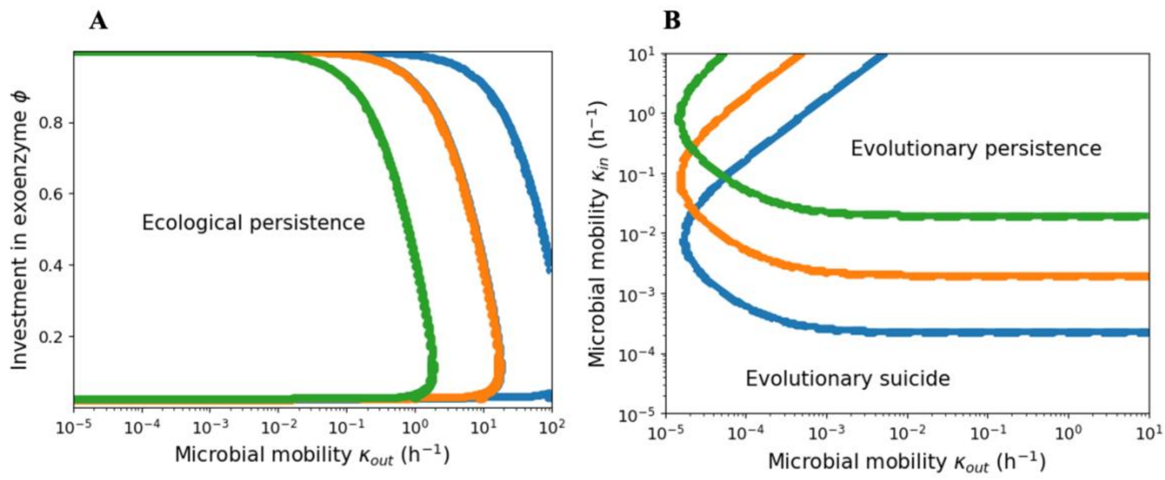


**Figure S1 | Zones of ecological and evolutionary persistence depending on microbial mobility and exoenzyme investment.** **(A)** In the zone to the left of the curves, ecological persistence of the microbial population is possible; in the zone to the right, the microbial population crashes. $\delta_m$= $10^{12}$ $m^{-3}$ and $\kappa_{in}$ = 0.1 for the green line, $\kappa_{in}$ = 1 for the orange line **and** $\kappa_{in}$ = 10 for the blue line. **(B)** In the top-right corner, evolutionary persistence of the microbial population is possible; in the bottom, cheaters take over the population until there is an evolutionary suicide. The blue line is for $\delta_m$= $10^{12}$ $m^{-3}$, the orange one for $\delta_m$= $10^{11}$ and the green one for $\delta_m$= $10^{10}$.

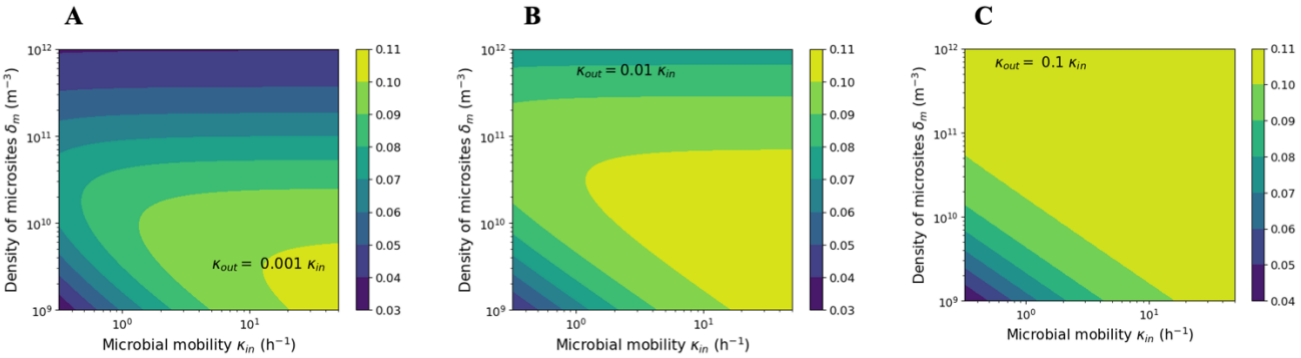


**Figure S2 | Evolutionary adapted investment in exoenzyme production, $\phi^*$, as a function of soil and microbial parameters at T = 20°C.** The adapted value is shown as a function of microsite density and microbial mobility, for **(A)** $\kappa_{out} = 0.001\ \kappa_{in}$; **(B)** $\kappa_{out} = 0.01\ \kappa_{in}$ and **(C)** $\kappa_{out} = 0.1\ \kappa_{in}$.

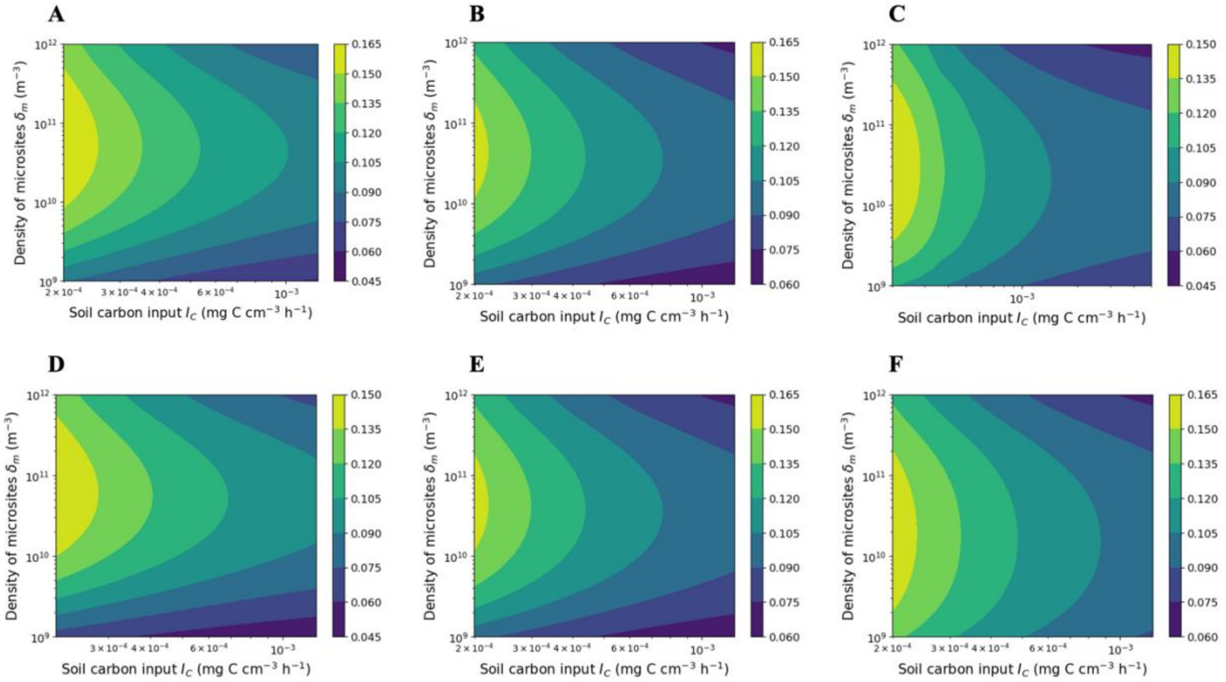


**Figure S3 | Evolutionary adapted investment in exoenzyme production, $\phi^*$, as a function of soil parameters.** The adapted value is shown as a function of microsite density and soil carbon input, for **(A)** T = 15°C and $\kappa_{in}$ = 1; **(B)** T = 20°C and $\kappa_{in}$ = 1; **(C)** T = 25°C and $\kappa_{in}$ = 1; **(D)** T = 20°C and $\kappa_{in}$ = 0.5; **(E)** T = 20°C and $\kappa_{in}$ = 1; **(F)** T = 20°C and $\kappa_{in}$ = 5.

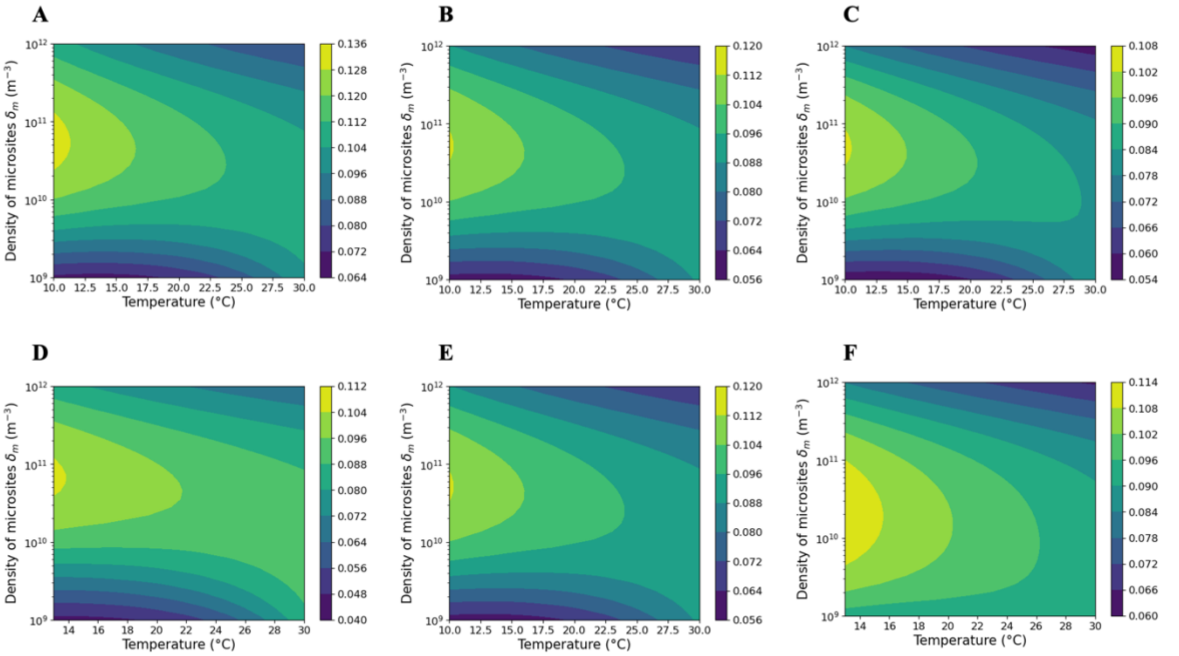


**Figure S4 | Evolutionary adapted investment in exoenzyme production, $\phi^*$, as a function of soil parameters and temperature.** The adapted value is shown as a function of microsite density and temperature, for **(A)** $I_C = 5 \times 10^{-4}$ and $\kappa_{in} = 1$; **(B)** $I_C = 10^{-3}$ and $\kappa_{in} = 1$; **(C)** $I_C = 2 \times 10^{-3}$ and $\kappa_{in} = 1$; **(D)** $I_C = 10^{-3}$ and $\kappa_{in} = 0.5$; **(E)** $I_C = 10^{-3}$ and $\kappa_{in} = 1$; **(F)** $I_C = 10^{-3}$ and $\kappa_{in} = 5$.

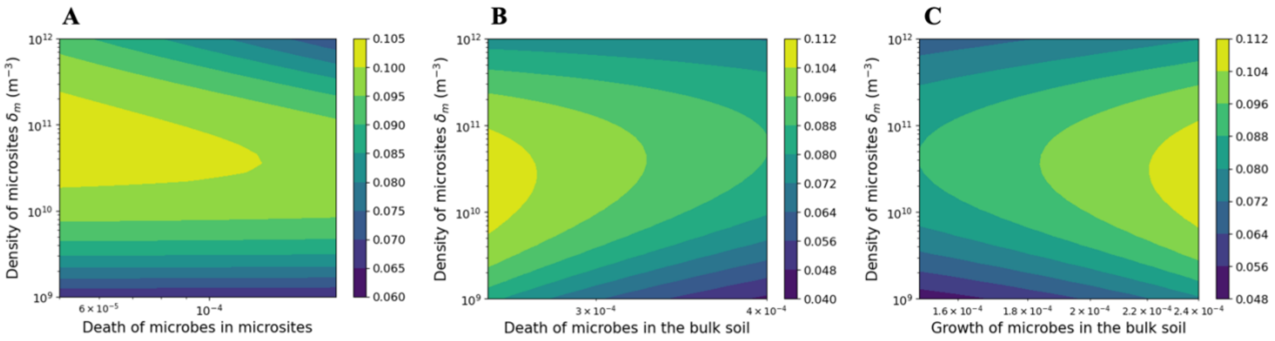


**Figure S5 | Evolutionary adapted investment in exoenzyme production, $\phi^*$, as a function of soil and microbial parameters at T = 20°C.** The adapted value is shown as a function of microsite density and **(A)** microbial death in microsites, **(B)** microbial death in the bulk soil and **(C)** microbial growth in the bulk soil.

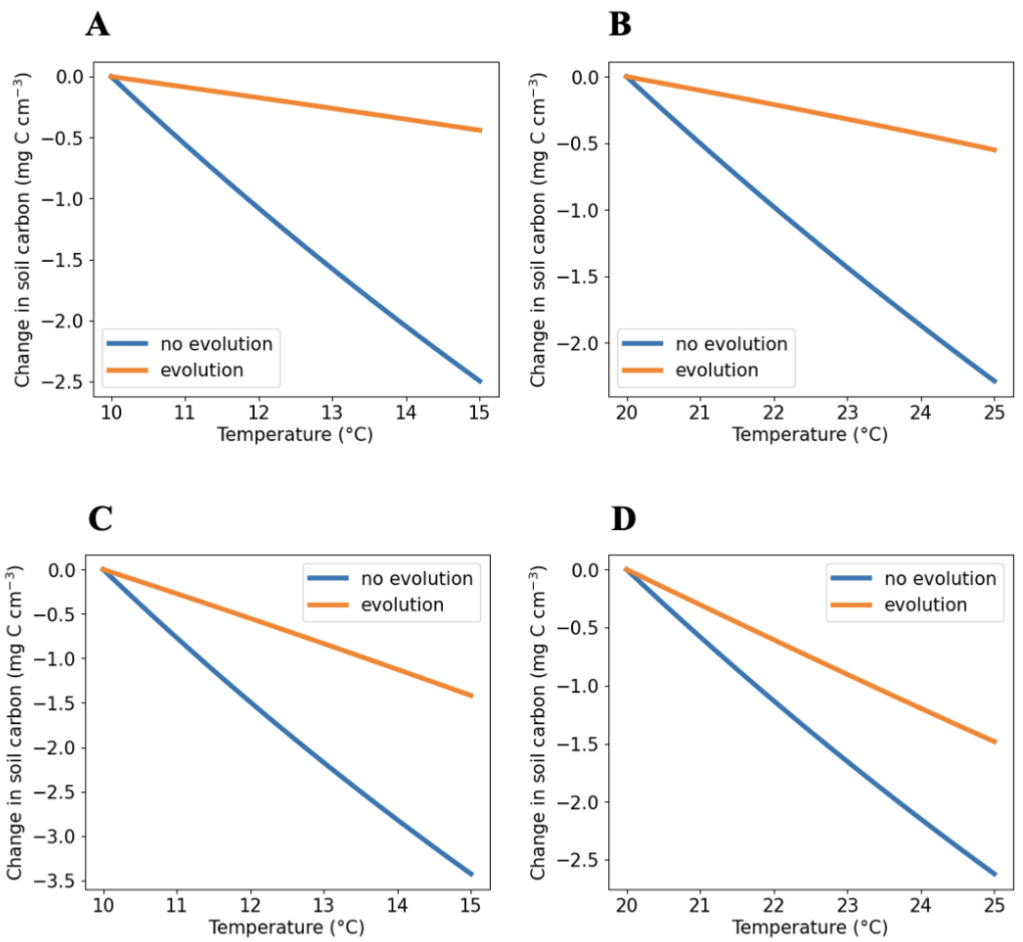


**Figure S6 | Soil carbon loss with and without evolutionary adaptation of microbial exoenzyme production to warming: influence of spatial structure parameters when microbial mortality depends on temperature. (A)** and **(B)** Microsite density $\delta_m = 10^{11}$ ($m^{-3}$). **(C)** and **(D)** Microsite density $\delta_m = 10^{9}$ ($m^{-3}$).
